# Residual Breast Cancer Cells Co-opt SOX5-driven Endochondral Ossification to Maintain Dormancy

**DOI:** 10.1101/2025.05.07.652632

**Authors:** Amulya Sreekumar, Eric Blankemeyer, Christopher J. Sterner, Tien-Chi Pan, Dhruv K. Pant, Sarah Acolatse, Hamza Turkistani, George K. Belka, Sean D. Carlin, Charles A. Assenmacher, Mark A. Sellmyer, David A. Mankoff, Lewis A. Chodosh

## Abstract

Recurrent breast cancer accounts for most disease-associated mortality and can develop decades after primary tumor therapy. Recurrences arise from residual tumor cells (RTCs) that can evade therapy in a dormant state, however the mechanisms are poorly understood. CRISPR-Cas9 screening identified the transcription factors SOX5/6 as functional regulators of tumor recurrence. Loss of SOX5 accelerated recurrence and promoted escape from dormancy. Remarkably, SOX5 drove dormant RTCs to adopt a cartilage-dependent bone development program, termed endochondral ossification, that was confirmed by [^18^F]NaF-PET imaging and reversed in recurrent tumors escaping dormancy. In patients, osteochondrogenic gene expression in primary breast cancers or residual disease post-neoadjuvant therapy predicted improved recurrence-free survival. These findings suggest that SOX5-dependent mesodermal transdifferentiation constitutes an adaptive mechanism that prevents recurrence by reinforcing tumor cell dormancy.

## Main Text

Despite significant advances in early detection and treatment, breast cancer remains the leading cause of cancer-related mortality in women worldwide (*1*). Most breast cancer-associated deaths result from therapy-refractory metastatic recurrences seeded by sub-clinical minimal residual disease (MRD) consisting of disseminated residual tumor cells (RTCs) that evaded treatment strategies targeting primary tumors (*2-4*). Consequently, depleting or eliminating MRD in patients via therapies that specifically target the mechanisms by which RTCs persist and escape dormancy has the potential to prevent tumor recurrence and its associated mortality (*5, 6*). However, such approaches are not yet standard clinical practice due to limited knowledge regarding how dormant RTCs survive and subsequently recur (*7*).

### In vivo CRISPR-Cas9 screening identifies *Sox5/6* as regulators of clonal enrichment

To identify candidate mechanisms regulating the switch from dormant residual disease to proliferating recurrent tumors, we analyzed data from a targeted in vivo CRISPR-Cas9 screen in *MMTV-rtTA;TetO-Her2/neu* (*MTB/TAN*)-derived cells (*8-10*), previously used to identify regulators of RTC survival (*11*) **(Fig. 1A)**. In this clinically validated *MTB/TAN* genetically engineered mouse (GEM) model (*11, 12*), doxycycline drives HER2 expression and primary tumor growth, following which withdrawal of doxycycline simulates neoadjuvant therapy by downregulating HER2, resulting in disease regression to a state of dormant MRD. Spontaneous, HER2-independent recurrences arise from MRD after variable periods of latency, as is observed in breast cancer patients.

**Fig. 1:**
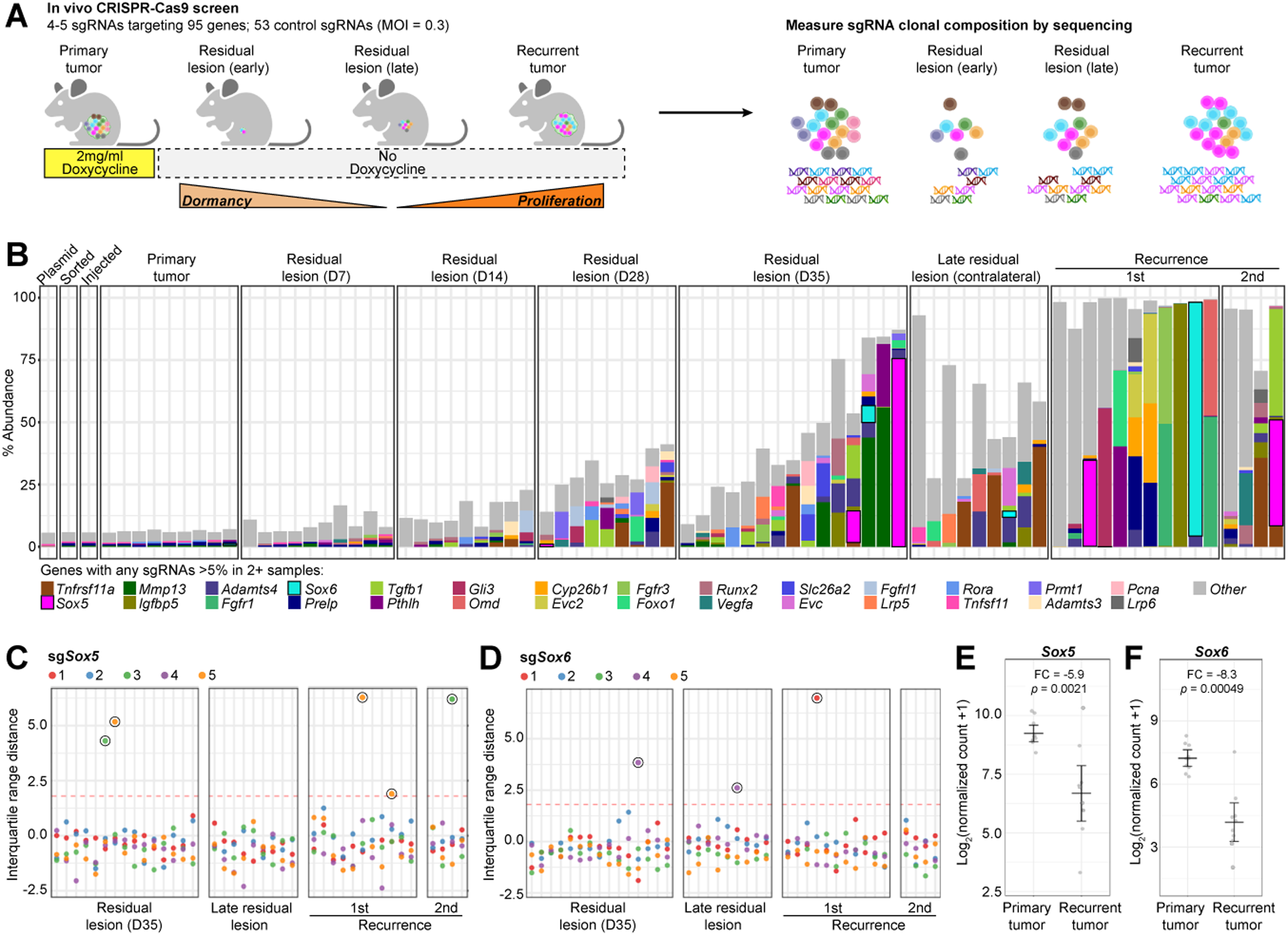
In vivo CRISPR-Cas9 screening identifies *Sox5*/*6* as regulators of clonal enrichment. **A.** In vivo CRISPR-Cas9 screen schematic using *MTB/TAN* cells. sgRNA composition measured in primary tumors, residual lesions, and recurrent tumors by next-generation sequencing. **B.** Stacked bar plot displays % abundance of top 10 sgRNAs in each sample. Colored bars indicate top 10 sgRNAs targeting genes that reached >5% in 2+ samples. Gray bar indicates other sgRNAs that did not meet this criterion. sg*Sox5* (fuchsia) and sg*Sox6* (cyan) outlined in black. **C.** Interquartile range distance of each of 5 sgRNAs targeting *Sox5* and **D.** *Sox6* in late residual lesions and recurrences. Red dotted line indicates the threshold of 1.8 used to call outliers. **E.** *Sox5* and **F.** *Sox6* gene expression in *MTB/TAN* primary tumors and spontaneous recurrences.

HER2-dependent Cas9 expressing tumor cells were transduced with a targeted library composed of ∼500 sgRNAs, resulting in a 0.2% abundance/sgRNA with similar representation across sgRNAs in the injected cell pool **(fig. S1A)**. Using a stringent cutoff of 5% abundance/sgRNA to identify clonally enriched sgRNAs, we observed a progressive enrichment of sgRNAs at later residual lesion time points and in recurrent disease **(Fig. 1B and fig. S1B)**. The enrichment of sgRNAs was particularly striking in recurrent tumors, which were generally clonal or oligoclonal and composed of a small number of sgRNAs **(Fig. 1B and fig. S1B)**. The marked and progressive skewing of sgRNA composition in late residual lesions and recurrent tumors was confirmed by Gini indices >0.90 (vs. 0.26 in the injected pool) (*11*).

To identify genes whose deletion might promote dormancy exit, we asked which sgRNAs were highly enriched at later stages of residual disease and recurrence. The top 10 genes targeted by sgRNAs that were present at >5% abundance in multiple samples were ranked as follows: *Tnfrsf11a*, *Sox5*, *Mmp13*, *Igfbp5*, *Adamts4*, *Fgfr1*, *Sox6*, *Prelp*, *Tgfb1*, and *Pthlh* **(Fig. 1B)**.

The enrichment in recurrent tumors of sgRNAs targeting either *Sox5* or *Sox6* – two members of the highly conserved SoxD group of the SRY-related HMG box (SOX) transcription factor family (*13*) was striking **(Fig. 1, C and D, and fig. S1, C and D)**. Consistent with a model in which loss of SOX5/6 transcription factors promotes clonal outgrowth from a dormant state, we also observed enrichment for sgRNAs targeting 3 downstream SOX5/6 targets included in our screen: *Pthlh*, *Runx2*, and *Tnfsf11* **(Fig. 1B)** (*14*). Moreover, comparing gene expression profiles from *MTB/TAN*-derived primary tumors and spontaneous recurrences identified a significant downregulation of *Sox5* (∼6-fold, *p* < 0.01) and *Sox6* (∼8-fold, *p* < 0.001) in recurrent tumors **(Fig. 1, E and F)**. Together, these data suggest that SOX5/6 downregulation promotes tumor recurrence.

### Loss of SOX5 promotes local and metastatic recurrence

To test the hypothesis that SOX5/6 loss promotes recurrence, we first transduced HER2*-* dependent-Cas9 cells with control sg*Rosa*-GFP or sg*Sox5-*GFP/sg*Sox6*-GFP. ICE analysis confirmed that sg*Sox5_5* and sg*Sox5_3* induced loss-of-function mutations in 91% or 83% of tumor cells, respectively, whereas sg*Sox6_1* and sg*Sox6_4* induced loss-of-function mutations in 83% or 45% of tumor cells, respectively **(fig. S2, A to D)**.

Having identified the most efficient sgRNAs targeting *Sox5* and *Sox6*, we performed an orthotopic recurrence-free survival assay to determine the impact of SOX5/6 deletion on median time to spontaneous recurrence in the mammary gland **(Fig. 2A)**. Because SOX5 and SOX6 are closely related paralogs with redundant functions (*13*), we also asked if deletion of both SOX5 and SOX6 results in a stronger phenotype. To assess the impact of targeting SOX5 and SOX6 singly or dually, HER2-dependent Cas9 cells were co-transduced with: i) sg*Rosa1* + sg*Rosa2*; ii) sg*Rosa1* + sg*Sox5_5*; iii) sg*Rosa1* + sg*Sox5_3*; iv) sg*Rosa2* + sg*Sox6_1*; or v) sg*Sox6_1* + sg*Sox5_5* and injected orthotopically into mice to drive doxycycline-dependent primary tumor outgrowth. While no differences in primary tumor formation or growth were observed upon loss of SOX5, loss of SOX6 marginally slowed primary tumor growth **(fig. S2E)**.

**Fig. 2:**
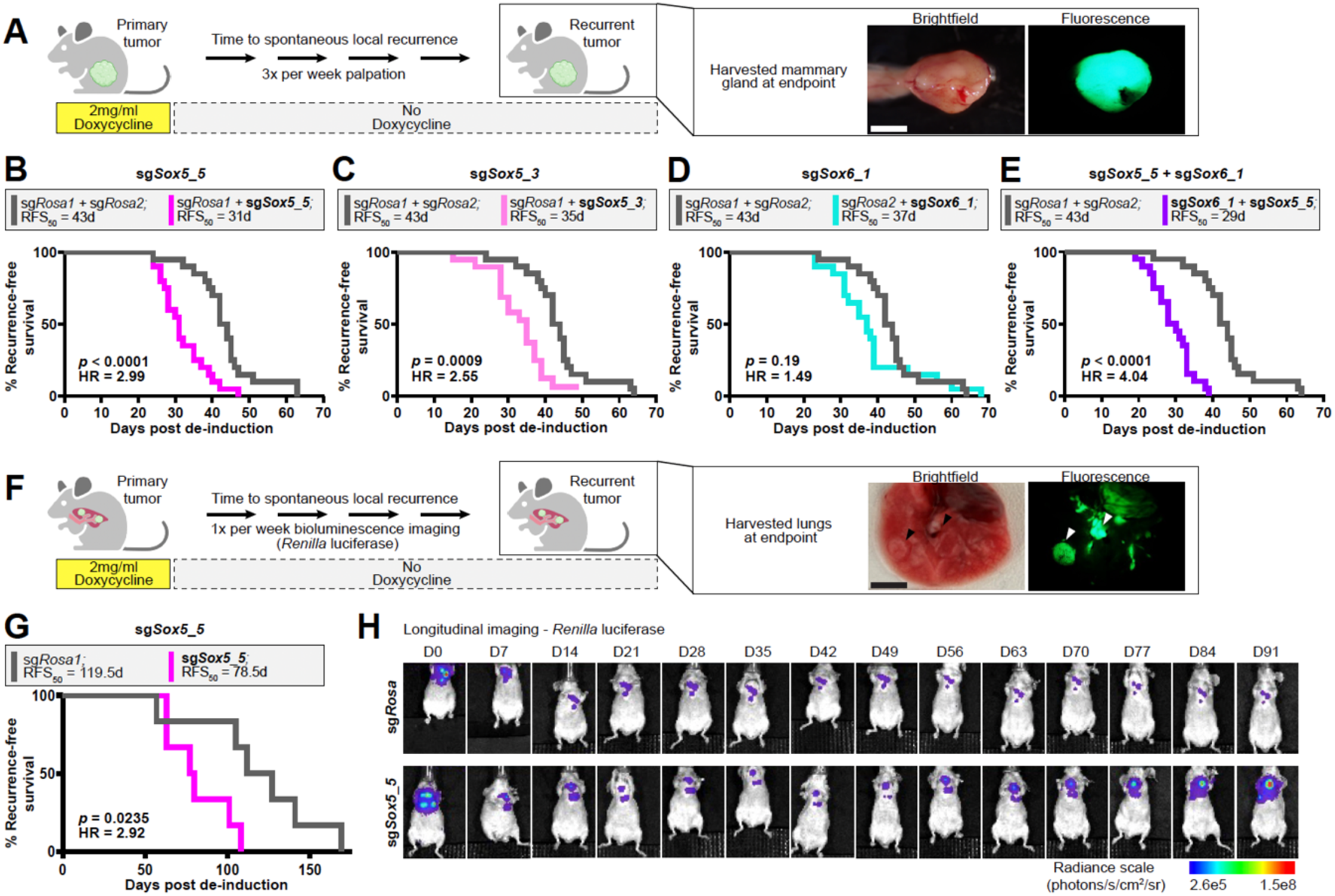
Loss of SOX5 promotes local and metastatic recurrence. **A.** Local recurrence-free survival schematic. Stereoscope images depict orthotopic mammary gland at endpoint. Scale bar = 5mm. **B-E.** Kaplan-Meier analysis of recurrence-free survival for *sgRosa1* + sgRosa2 (gray), **B.** *sgRosa1* + sgSox5_5 (fuchsia), **C.** *sgRosa1* + sgSox5_3 (light pink), **D.** sgRosa2 + sgSox6_ *1* (cyan), and **E.** sgRosa2 + sgSox6_ *1* + sgSox5_5 (purple) groups. n = 20 mice/group. RFS_50_ = median time-to-recurrence. **F.** Experimental metastasis recurrence-free survival schematic. Stereoscope images depict lungs at endpoint. Arrowheads indicate visual recurrent tumors in the lungs, which express GFP. Scale bar= 5mm. **G.** Kaplan-Meier analysis of experimental metastasis recurrence-free survival for *sgRosa1* (gray) and sgSox5_5 (fuchsia). n = 6 mice/group. RFS_50_ = median time-to-recurrence. **H.** Bioluminescence imaging for peak *Renilla* luciferase activity following coelenterazine injection.

As predicted from our CRISPR-Cas9 screen, deletion of *Sox5* by sg*Sox5_5* or sg*Sox5_3* strongly accelerated time-to-recurrence (*p* < 0.0001, HR = 2.99; and *p* = 0.0009, HR = 2.55, respectively) **(Fig. 2, B and C)**. Additionally, consistent with the lower rank of *Sox6* in the CRISPR-Cas9 screen, sg*Sox6* modestly accelerated time-to-recurrence (HR = 1.49), although this did not reach statistical significance **(Fig. 2D)**. Notably, the strongest acceleration of recurrence was observed for dual knockout of *Sox5* and *Sox6* (*p* < 0.0001, HR = 4.04) **(Fig. 2E)**. Moreover, the difference between dual *Sox5/6* knockout **(Fig. 2E)** and *Sox5* knockout alone **(Fig. 2B)** was not significant, suggesting that the dominant effect on recurrence is driven by SOX5. No significant differences were observed in the growth rates of recurrent tumors across groups **(fig. S2F).**

Next, we asked whether loss of SOX5 also promotes tumor recurrence at metastatic locations such as the lung, which is a major site of metastatic recurrence in breast cancer patients (*15*). HER2-dependent cells harboring sg*Rosa1* or sg*Sox5_5* and constitutively expressing *Renilla* luciferase were injected into mice maintained on doxycycline via tail vein injection. Oncogene-dependent tumor growth was monitored weekly using Firefly luciferase, expression of which is also driven by doxycycline along with the HER2 in *MTB/TAN* tumor cells (*9*) **(Fig. 2F)**. When Firefly luciferase emission exceeded 10^9^ photons/s in the mouse **(fig. S2, G and H)**, doxycycline was withdrawn and *Renilla* luciferase activity was monitored weekly. As anticipated, *Renilla* luciferase activity initially decayed ∼50-fold as a consequence of tumor regression induced by HER2 downregulation upon doxycycline withdrawal. *Renilla* luciferase activity was detected at consistently low levels for several weeks before spontaneous increase in signal attributable to recurrent tumor growth in the lung **(fig. S2, I and J)**. Recurrences developed significantly faster in the sg*Sox5_5* group compared to sg*Rosa* (*p* = 0.0235, HR = 2.92) **(Fig. 2, G and H)**, suggesting that SOX5 regulates breast cancer latency at metastatic as well as primary tumor sites.

### Loss of SOX5 impairs maintenance of RTC dormancy

In light of our observations that SOX5 loss promotes tumor recurrence, and that sg*Sox5*-containing tumor cells were clonally enriched at later residual lesion time points prior to recurrence, we hypothesized that SOX5 might enforce a dormant state in RTCs. To address this, we first assessed *Sox5* expression and predicted activity in dormant RTCs. *Sox5* was highly expressed in HER2-dependent cells in vitro and further upregulated at multiple dormancy time points **(Fig. 3A)**. Furthermore, a predicted SOX5 regulon composed of 19 genes (*14*) was markedly upregulated in tumor cells beginning at Day 7 (D7) following doxycycline withdrawal – the first time point at which the great majority of cells reside in dormant state (*11*) – and was further upregulated at D14 and D28 as cells progressed deeper into dormancy **(Fig. 3B)**. Moreover, SOX5 activity was reversibly downregulated upon cell cycle reentry induced by doxycycline re-addition, suggesting that its transcriptional activation is dormancy specific.

**Fig. 3:**
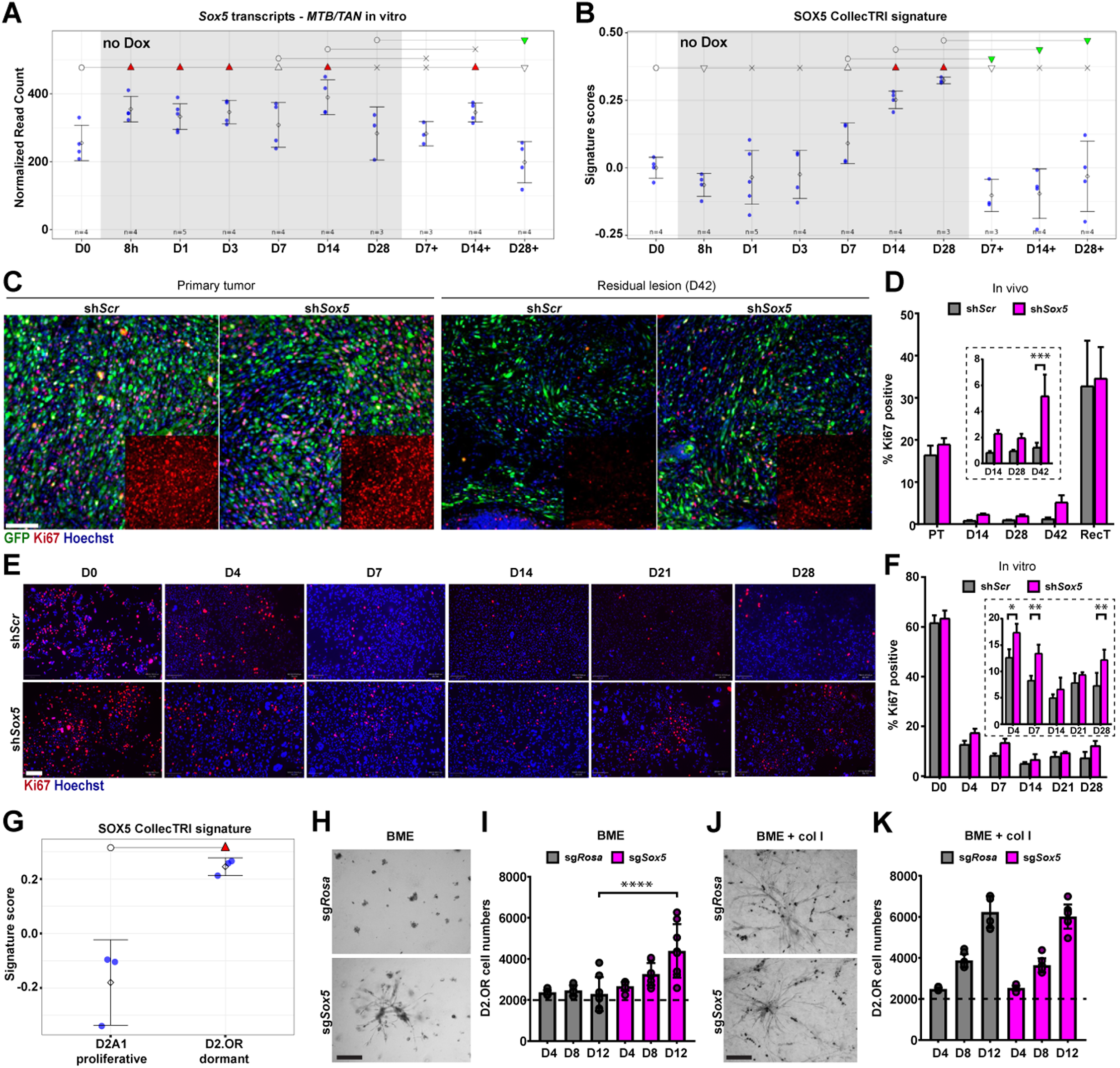
Loss of SOX5 impairs maintenance of RTC dormancy. **A.** *Sox5 MTB/TAN* IVD temporal gene expression profile and **B.** CollecTRI SOX5 regulon signature temporal expression profile. Circle = baseline, x = non-significant, empty triangles = trending significance 0.05 < *p* < 0.1, filled triangles = *p* < 0.05, inverted triangles = decreased signature score, upright triangles = increased signature score. **C.** Immunofluorescence and **D.** quantification for Ki67 (red) in tumor cells (green) in sh*Scr* (gray) and sh*Sox5* (dark green) groups in vivo. Quantification is represented as mean ± SD. n = 4 PT and RT/group, n = 6 RL/group. \**p* < 0.05, \*\*\**p* < 0.001. Scale bar = 100µm. **E.** Immunofluorescence and **F.** quantification for Ki67 (red) in tumor cells (green) in sh*Scr* (gray) and sh*Sox5* (fuchsia) groups in vitro. Quantification is represented as mean ± SD. n = 3 replicates/group. \**p* < 0.05, \*\**p* < 0.01. Scale bar = 200µm. **G.** CollecTRI SOX5 regulon signature in D2.OR/D2A1 cells grown in 3D (basement membrane extract (BME)). Circle = baseline, filled triangles = *p* < 0.05, upright triangles = increased signature score. **H-K.** Brightfield images of sg*Rosa* and sg*Sox5* D2.OR cells grown in 3D on **H.** basement membrane extract (BME) or **J.** BME + collagen I (col I) and viable cell numbers in **I.** BME or **K.** BME + col I measured at D4, D8, and D12 time points. Dotted line indicates cell number at D0. Scale bar = 200µm. Data are represented as mean ± SD. n = 8 replicates/group. \*\*\*\**p* < 0.0001.

To determine whether SOX5 is required to maintain RTC dormancy, we used HER2-dependent cells transduced with a sh*Sox5* hairpin that demonstrated ∼80% knockdown of *Sox5* **(fig. S3A)** and recapitulated the acceleration of tumor recurrence observed for sgRNAs targeting *Sox5* in HER2-dependent Cas9 cells without altering recurrent tumor growth rates **(fig. S3, B and C)**. This system enabled examination of a more protracted latency period with median time-to-recurrence of 60 days **(fig. S3B)**.

To determine the cellular mechanisms underlying the accelerated recurrence observed upon loss of SOX5, we performed immunofluorescence for Ki67 on sh*Scr*-GFP or sh*Sox5*-GFP primary tumors (PT), residual lesions following 14, 28, or 42 days of HER2 downregulation (D14, D28, D42), and recurrent tumors (RecT). While no differences in %Ki67-positive cells were observed in primary tumors, we observed increased levels of RTC proliferation in residual lesions that became most pronounced by D42 **(Fig. 3, C and D)**. These data suggest that SOX5 loss results in impaired maintenance of the dormant state.

To determine whether the increased RTC proliferation observed at dormancy time points as a consequence of SOX5 loss is tumor-cell autonomous, we performed an in vitro dormancy assay in HER2-dependent cells (*11*). Knockdown, as well deletion, of *Sox5* in vitro resulted in progressive increases in the number of viable cells **(fig. S3, D and E)** that were accompanied by increased RTC proliferation as measured by Ki67 immunofluorescence **(Fig. 3, E and F, and fig. S3, F and G)**. These data suggest that SOX5 loss impairs dormancy maintenance in a cell autonomous manner.

To determine whether SOX5-induced RTC dormancy is a more general mechanism of adaptation to stress, we extended our studies to a paradigm of microenvironment-induced dormancy. To this end, we used the D2.OR-D2A1 paired tumor cell line model, which proliferate comparably when grown on plastic in 2-dimensions (2D) but exhibit dormant (D2.OR) or proliferative (D2A1) behaviors, respectively, when plated in 3-dimensions (3D) in basement membrane extract (BME). Using previously published gene expression data (*16*), we first determined whether the SOX5 regulon described above was selectively enriched in dormant cells. Indeed, dormant D2.OR cells exhibited significant upregulation of SOX5 activity relative to proliferative D2A1 cells when grown in 3D **(Fig. 3G)**, but not in 2D (data not shown).

To assess whether SOX5 is required for microenvironment-induced dormancy, we used D2.OR cells that are dormant when grown on BME but proliferate when grown on BME supplemented with collagen I (col I). D2.OR cells were nucleofected with Cas9:sgRNA ribonucleoprotein complex to generate sg*Rosa* and sg*Sox5* D2.OR cells. After confirming efficient knockout **(fig. S3, H and I)**, sg*Rosa* and sg*Sox5* D2.OR cells were plated on BME or BME + col I matrices.

As expected, the number of sg*Rosa* D2.OR cells were unchanged over a 12-day period. In contrast, the number of sg*Sox5* D2.OR cells increased progressively over this time frame and was accompanied by signature proliferative outgrowths **(Fig. 3, H and I)**. Conversely, no differences between sg*Rosa* and sg*Sox5* genotypes were observed when D2.OR cells were grown on BME + col I **(Fig. 3, J and K)**. These data indicate that SOX5 is required for maintaining the dormant state of RTCs independently of whether dormancy is induced by targeted therapy or by microenvironmental cues.

### RTCs upregulate an endochondral ossification program

Our data thus far were consistent with a model wherein SOX5 is required for promoting RTC dormancy, thereby delaying recurrence. To investigate the mechanism by which SOX5 might promote dormancy, we turned to known physiological functions of SOX5. Heterozygous mutations in *SOX5* are causally linked to a neurodevelopmental disorder, Lamb-Shaffer syndrome, which is characterized by intellectual disability, speech and locomotor retardation, and skeletal abnormalities (*17*). Indeed, SOX5 has a well characterized role in skeletal development where, together with SOX6 and SOX9, it coordinates and promotes a program of chondrogenesis-dependent bone formation, termed endochondral ossification (*18-20*). Therefore, we asked if similar SOX5-dependent mechanisms might be operative in dormant MRD.

As a first step, we performed gene ontology analysis on a core RTC signature derived from integrating in vitro and in vivo-derived gene expression from the *MTB/TAN* model (*11*) and asked which biological processes were enriched in dormant RTCs. The top 25 pathways enriched in dormant *MTB/TAN* RTCs included *Extracellular matrix organization*, *Cell adhesion*, *Cell migration*, and *Cell differentiation*. Strikingly, *Ossification* was significantly enriched in dormant RTCs compared to their proliferative counterparts and was the 4^th^-highest pathway as assessed by magnitude of enrichment **(Fig. 4A)**.

**Fig. 4:**
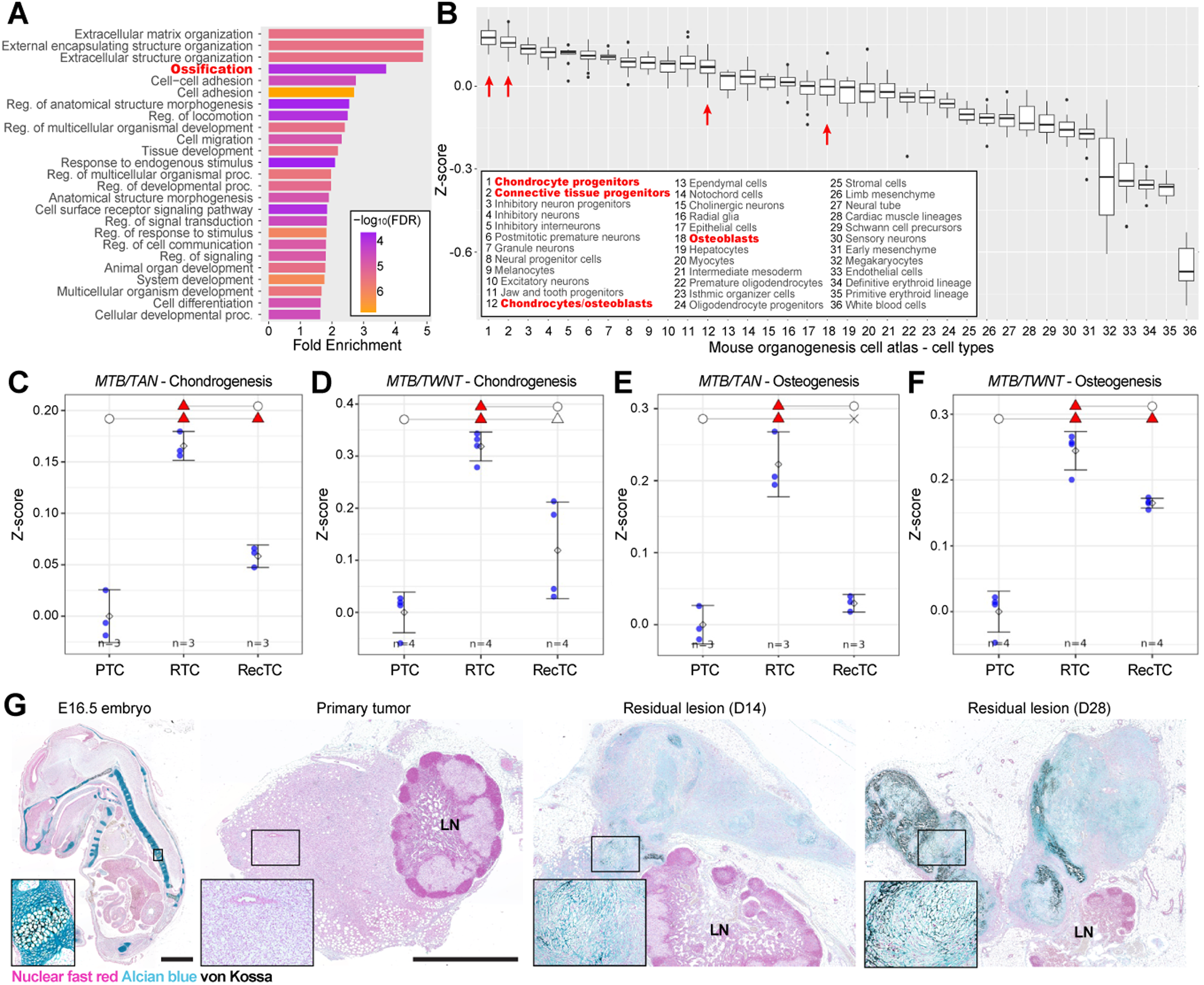
RTCs upregulate an endochondral ossification program. **A.** Top 25 gene ontology - biological process terms for dormancy upregulated genes. Ossification highlighted in red. **B.** Tabula muris analysis of dormancy upregulated genes. Chondrocyte and osteoblast-related cell types are highlighted in red. **C-F.** Application of the chondrogenesis and osteogenesis signatures on primary tumor cells (PTCs), residual tumor cells (RTCs), and recurrent tumor cells (RecTCs) derived from **C, E.** *MTB/TAN* and **D, F.** *MTB/TWNT* models. Circle = baseline, x = non-significant, empty triangles = trending significance 0.05 < *p* < 0.1, filled triangles = *p* < 0.05. **G.** Alcian blue (blue) and von Kossa (black) staining to identify cartilage- and bone-like areas, respectively. LN = lymph node. Scale bar = 2mm.

To identify which specific aspects of ossification were exhibited by dormant RTCs, we applied the core RTC signature to a mouse organogenesis cell atlas that profiled ∼2 million single cells from mouse embryos between E9.5 and E13.5 (*21*). This revealed that the core RTC signature was most highly enriched in chondrocyte progenitors and connective tissue progenitors among the 36 cell types annotated in this mouse organogenesis cell atlas, with chondrocytes and osteoblasts also demonstrating enrichment **(Fig. 4B)**.

The co-occurrence of chondrocyte differentiation with ossification motivated us to ask whether RTCs might exhibit evidence of endochondral ossification. To this end, we applied chondrogenesis- and osteogenesis-associated signatures derived from the in vitro differentiation of fibroblasts (*22*) to gene expression profiles of sorted primary tumor cells (PTCs), dormant residual tumor cells (RTCs), and recurrent tumor cells (RecTCs) from the *MTB/TAN* and *MTB/TWNT* (*MMTV-rtTA;TetO-Wnt1*) mouse models in vivo (*12, 23*). In both the HER2 and WNT1 GEM models, chondrogenic differentiation **(Fig. 4, C and D)** and osteogenic differentiation **(Fig. 4, E and F)** signatures were each specifically and highly enriched in dormant RTCs compared to PTCs and were downregulated in RecTCs. Furthermore, both chondrogenic and osteogenic differentiation signatures were progressively enriched during dormancy in vitro following HER2 downregulation, suggesting that engagement of chondrogenic and osteogenic differentiation programs in dormant RTCs is tumor cell-autonomous **(fig. S4, A and B)**.

To confirm the prediction based on gene expression signatures that RTCs in MRD undergo osteochondrogenic transdifferentiation, we performed histological staining using Alcian blue and von Kossa. Alcian blue stains acidic polysaccharides that are abundantly synthesized by chondrocytes, whereas von Kossa employs a silver nitrate solution to replace calcium ions with silver ions in a light-catalyzed reaction to detect bone calcification. This combination of stains can be used to clearly demarcate skeletal elements (*20*) as observed in a positive control section from an E16.5 mouse embryo **(Fig. 4G)**. As predicted from gene expression profiling, Alcian blue and von Kossa staining of primary tumors and dormant residual lesions from the *MTB/TAN* model revealed the selective and pronounced enrichment of osteochondrogenic differentiation in D14 and D28 residual lesions compared to primary tumors **(Fig. 4G)** within the regions occupied by GFP+ RTCs **(fig. S4C)**. Together, these data suggest that dormant epithelial RTCs undergo transdifferentiation to a mesodermal fate.

### Dormant RTCs temporally upregulate ossification in a SOX5-dependent manner

To study the kinetics of ossification during tumor progression, we performed longitudinal imaging of mice injected orthotopically into the #4 inguinal mammary fat pad with HER2-dependent cells. Computed tomographic (CT) imaging was performed in concert with positron emission tomography (PET) using [^18^F]NaF, a radiopharmaceutical targeting endochondral and bony matrix mineralization that is used clinically for detecting benign and malignant bone lesions (*24-26*). [^18^F]NaF tracer uptake overlaid well with CT signal in the skeleton and was rapidly cleared through the urine with expected extra-skeletal signal in the urinary tract, including the bladder **(Fig. 5A)**. This approach revealed a temporal increase in extra-skeletal CT attenuation and PET tracer uptake at the site of the residual lesions **(Fig. 5B**, *white arrowheads*), consistent with ossification.

**Fig. 5:**
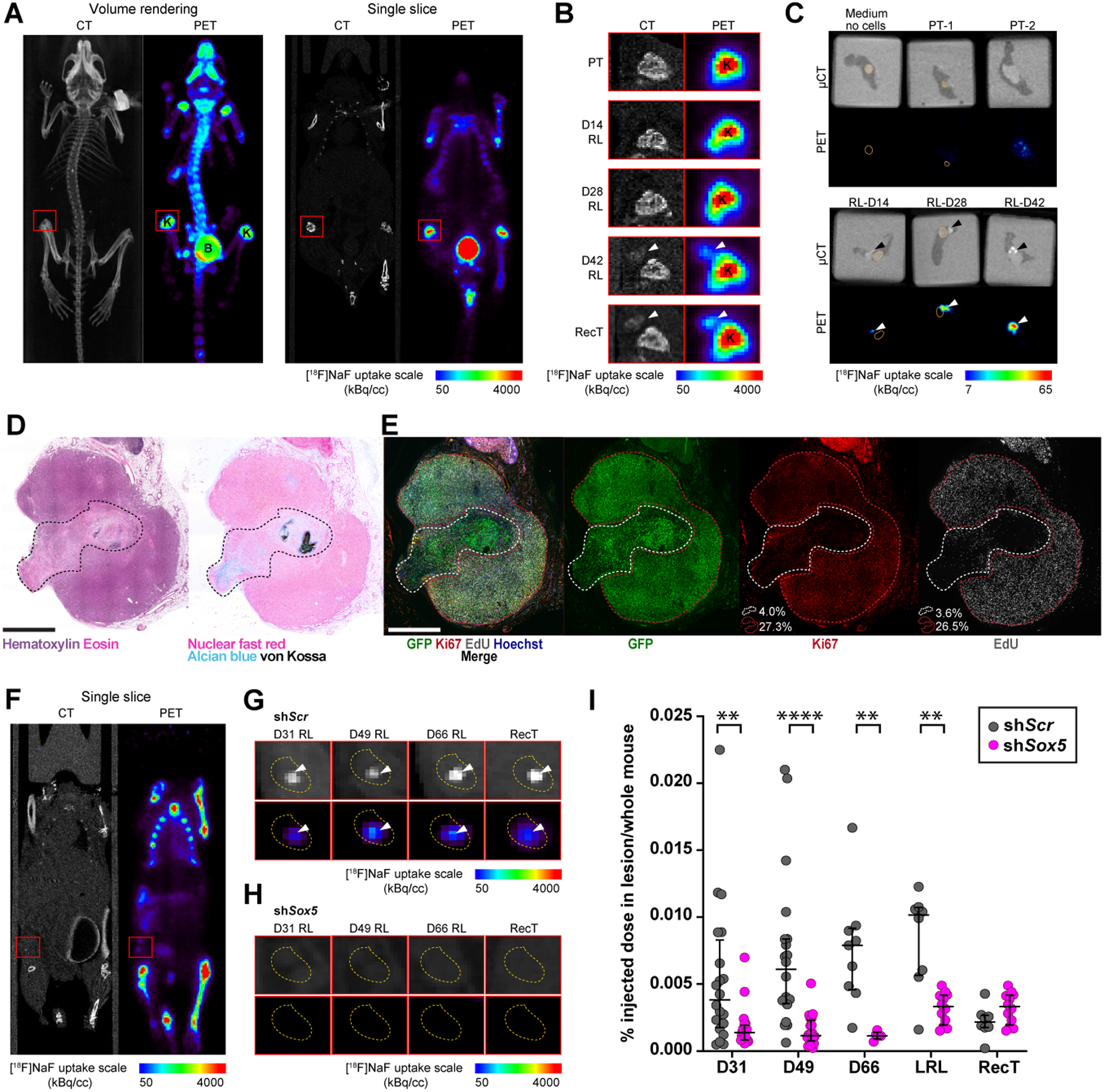
Dormant RTCs temporally upregulate ossification in a SOX5-dependent manner. **A.** Whole animal imaging by CT and PET 60 minutes after injection of [^18^F]NaF and **B.** zoomed-in view of signal in the orthotopic inguinal #4 mammary glands (red boxes). White arrowheads indicate calcified areas outside the knee joint. K = knee joint, B = bladder. **C.** µCT and PET 75 minutes after injection of [^18^F]NaF. Arrowheads indicate calcified areas in excised mammary glands. Orange dotted lines indicate lymph nodes. **D.** Serial sections of a recurrent tumor with a calcified lesion as identified by Hematoxylin and Eosin staining (panel 1) as well as Alcian blue (blue) and von Kossa (black) staining to identify cartilage- and bone-like areas, respectively (panel 2). Black dotted line indicates eosinophilic/calcified region. **E.** Immunofluorescence for GFP+ tumor cells (green), Ki67 (red), 24h EdU uptake (gray) in calcified ares (white dotted line) and non-calcified recurrent outgrowth (red dotted line). Percentage of tumor cells positive for Ki67 and EdU in each of these areas is indicated in the Ki67 and EdU staining panel. Scale bar = 2mm. **F.** Whole animal imaging by CT and PET 60 minutes after injection of [^18^F]NaF. Red boxes depict signal in the orthotopic inguinal #4 mammary glands. **G.** Yellow dotted lines indicate the boundaries of the orthotopic inguinal #4 mammary fat pad identified from whole mouse imaging of sh*Scr* and **H.** sh*Sox5* lesions. White arrowheads indicate calcified areas within the orthotopic inguinal #4 mammary glands. **I.** Quantification of the percent injected dose of [^18^F]NaF in the residual lesion normalized to whole mouse uptake in longitudinal imaging of n = 10 mice/group injected bilaterally in orthotopic inguinal #4 mammary glands. Quantification is represented as median ± interquartile range.\*\**p* < 0.01, \*\*\*\**p* < 0.0001.

Due to small lesion size and limited [^18^F]NaF spatial resolution in this whole animal imaging experiment, it was challenging to resolve tracer signal in the residual tumor lesion in the mammary gland from the nearby, more intense signal corresponding to the knee joint. Therefore, to further examine temporal patterns of ossification in residual lesions, we excised mammary glands bearing primary tumors or residual lesions and performed µCT and [^18^F]NaF-PET imaging ex vivo. This approach identified focal regions of calcification in tumors that were detectable by both CT and [^18^F]NaF tracer uptake in residual lesions, but not in PTs or mock-injected control glands **(Fig. 5C**, *arrowheads*). Notably, [^18^F]NaF tracer uptake progressively increased in dormant residual lesions in a time-dependent manner. Thus, [^18^F]NaF-PET/CT imaging constitutes a tractable approach to measuring and quantifying dynamic changes in calcification during cancer progression.

Although the highest levels of [^18^F]NaF-PET tracer uptake were observed in recurrent tumor samples **(fig. S5A)**, closer inspection revealed that in each case the visualized calcified focus was spatially distinct from the adjacent growing recurrent tumors **(fig. S5B)**. To characterize this spatial relationship in greater detail, we performed hematoxylin and eosin staining on sections and confirmed the presence of a visually distinct eosinophilic area embedded within the recurrent tumor mass **(Fig. 5D)**. Alcian blue/von Kossa staining of serial sections confirmed high levels of staining within this eosinophilic area **(Fig. 5D)**, consistent with our prior results and suggesting that it corresponded to the dormant residual lesion.

To directly address whether the eosinophilic, calcified focus embedded within the recurrent tumor does represent the dormant residual lesion, we performed EdU labeling (24h EdU uptake) and immunofluorescence for Ki67 to detect proliferating cells, as well as immunofluorescence for GFP to detect tumor cells, on serial sections. This confirmed that tumor cells within the eosinophilic, calcified region displayed markedly lower Ki67 and EdU positivity (∼4%) compared to the surrounding proliferating recurrent tumor (∼27%) **(Fig. 5E)**. These findings indicate that recurrent tumor samples are composed of two distinct entities: a late residual lesion that is calcified and dormant, and an adjacent recurrent tumor outgrowth that is proliferative and not calcified. More broadly, these data illustrate that ossification is associated with dormant residual lesions, as predicted from gene expression data **(Fig. 4, C to F and fig. S4, A and B).**

The above findings suggested that dormant, calcified residual lesions can give rise to adjacent actively proliferating recurrent tumors. Consistent with this, we also observed calcification in lung metastatic lesions in sg*Rosa* controls examined at recurrence endpoints **(Fig. 2F)** at the boundaries of non-calcified, proliferative outgrowths **(fig. S5C)**. Calcified areas in the lung existed around GFP+ tumor cells and demonstrated an inverse relationship with 24h EdU uptake **(fig. S5D)** similar to that observed for local residual lesions and recurrent tumors.

To test the hypothesis that SOX5 is required for the endochondral ossification program in dormant RTCs, we injected HER2-dependent cells harboring sh*Scr* control or sh*Sox5* hairpins and assessed dynamic calcification over time by longitudinal [^18^F]NaF-PET/CT imaging. In this experiment, we repositioned mice during imaging to spatially resolve signal in the orthotopic tumor site (*red boxes*) from uptake in the knee joint **(Fig. 5F)**. Using this approach, we were able to clearly detect calcification within residual lesions by CT and by [^18^F]NaF-PET, which showed increasing tracer uptake over time in dormant residual lesions **(Fig. 5G and fig. S5E)**.

In contrast to control lesions, calcification was markedly impaired in sh*Sox5* residual lesions across time points as visualized by both CT and [^18^F]NaF-PET tracer uptake **(Fig. 5H and fig. S5F)**. The SOX5-dependent difference was confirmed by quantification in longitudinally imaged mice wherein sh*Scr* residual lesions exhibited progressive increases in [^18^F]NaF-PET tracer uptake that were largely abrogated in sh*Sox5* residual lesions **(Fig. 5I)**. This association held true in late residual lesions adjacent to recurrent tumors, whereas recurrent tumors themselves exhibited low [^18^F]NaF tracer uptake **(Fig. 5I)**. The impairment of ossification in sh*Sox5* residual lesions was confirmed in a subset of orthotopic mammary glands that were excised for ex vivo [^18^F]NaF-PET/µCT imaging **(fig. S5, G and H)**. Together, these data indicate that depletion of the chondrogenic transcription factor SOX5 impairs endochondral ossification in dormant residual disease.

### Osteochondrogenic gene expression is enriched following neoadjuvant therapy in breast cancer patients and is associated with favorable prognosis

The above data provide evidence that dormant RTCs upregulate a SOX5-mediated endochondral ossification program. Because the chondrogenesis and osteogenesis gene expression signatures investigated above were found to provide reasonable proxies for endochondral ossification in dormant MRD **(Fig. 4, C to G)**, we applied these signatures to gene expression data from >4000 early-stage primary breast cancers with known recurrence-free survival data (*11, 12*). Consistent with our findings that osteochondrogenic differentiation promotes dormancy in residual disease, we observed that enrichment of chondrogenesis and osteogenesis signature in primary breast cancers was associated with markedly improved recurrence-free survival (*p* = 3.3e-6, HR = 0.58; and *p* = 3.0e-14, HR = 0.48, respectively) **(Fig. 6, A and B)**. Additionally, we found that chondrogenesis and osteogenesis signatures were strongly enriched in METABRIC integrative clusters containing breast cancers that did not relapse within 20 years of follow up, when compared to integrative clusters for early relapse (<5y) or late relapse (>5y) **(Fig. 6, C and D)** (*27*).

**Fig. 6:**
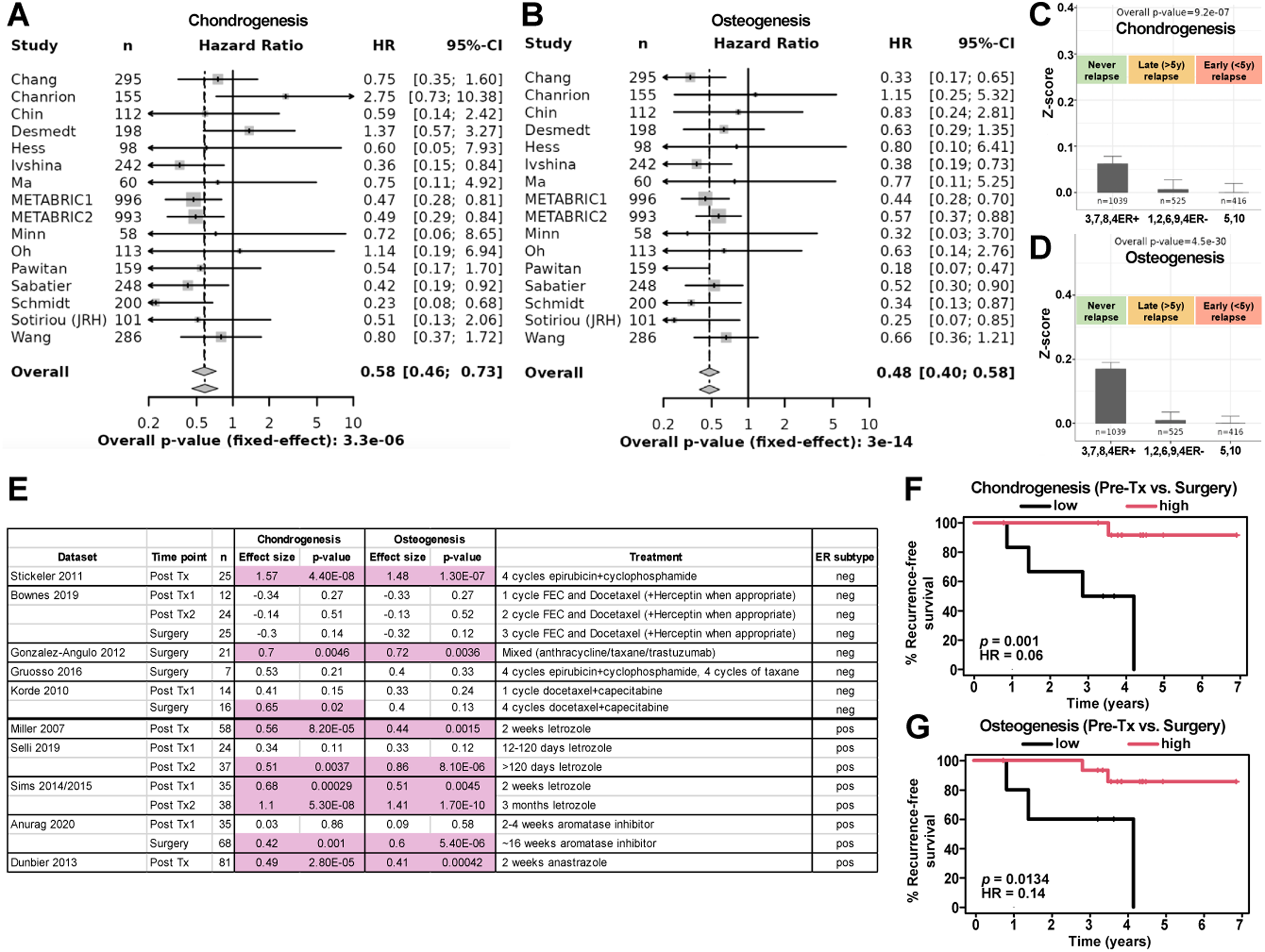
Osteochondrogenic gene expression is enriched following therapy and is associated with favorable patient prognosis. **A.** Forest plot representation of hazard ratios (HR) and 95% confidence intervals (CI) as a function of the chondrogenesis signature or **B.** osteogenesis signature in breast cancer patients recurring 10 years after initial treatment. Dashed lines depict the shift in HR across 16 human datasets. **C.** Application of the chondrogenesis and **D.** osteogenesis signatures on METABRIC integrative clusters grouped by patients who never relapse, demonstrate late relapse (>5 years after diagnosis), or demonstrate early relapse (<5 years after diagnosis). **E.** Chondrogenesis and osteogenesis signature enrichment following neoadjuvant therapies vs. pre-treatment (pre-tx) samples across 10 patient datasets measured by effect size (mean of pairwise difference in signature scores). Significant effects are highlighted in pink. **F, G.** Kaplan-Meier analysis of recurrence-free survival as a function of enrichment of **F.** chondrogenesis and **G.** osteogenesis signatures following neoadjuvant therapies vs. pre-tx in 21 patients.

Finally, we asked if chondrogenesis and osteogenesis signatures are enriched following neoadjuvant therapy as an adaptive response. We analyzed 10 publicly available datasets containing paired pre- and post-neoadjuvant therapy tissues samples in patients treated with either chemotherapy or endocrine therapy and encompassing both estrogen receptor (ER)-negative and ER+ disease (*28-38*). This revealed significant enrichment of the chondrogenesis signature in 8 of the 10 datasets examined, with enrichment of the osteogenesis signature observed in 7 of 10 datasets **(Fig. 6E)**.

Strikingly, in a subset of patients with post-neoadjuvant therapy gene expression data for whom recurrence outcomes were available (*39*), we found that enrichment of chondrogenic and osteogenic gene expression signatures were each associated with significantly improved recurrence-free survival (*p* = 0.001, HR = 0.06; and *p* = 0.0134, HR = 0.14, respectively) **(Fig. 6, F and G)**. These data provide evidence that osteochondrogenic differentiation of breast cancer cells in patients is associated with indolent disease and may represent a means of stratifying breast cancer patients who are less likely to develop recurrent tumors.

## Discussion

Up to 30% of early-stage breast cancer patients will ultimately develop lethal recurrent tumors following primary tumor therapy, principally due to the persistence of RTCs at local and distant sites (*40*). Since no clinically approved test currently exists to detect RTCs, virtually all early-stage patients are formally at risk for recurrence, resulting in uncertainty and decreased quality of life for patients and limited information for their physicians.

Long-standing observations that breast cancers may recur decades after initial treatment (*3, 4, 41*) led to the hypothesis that therapy-resistant RTCs may persist in their host in a dormant state. Unfortunately, the molecular mechanisms regulating tumor cell dormancy have remained elusive, thus precluding the identification of breast cancer patients at highest risk of recurrence and preventing the development of therapeutic approaches to specifically target and eliminate this critical reservoir of residual disease (*7, 42*). Here we provide evidence that the transcription factor SOX5 plays a pivotal role in suppressing tumor recurrence by virtue of its ability to enforce dormancy by promoting RTC transdifferentiation to co-opt an endochondral ossification program typically executed by mesodermal cells.

The transdifferentiation of ectodermally derived mammary epithelial cells towards mesodermal chondrogenic and osteogenic lineages as a mechanism driving dormancy is an unexpected phenomenon. Notably, multiple orthogonal findings support this claim and indicate that key regulators of endochondral bone development are selectively activated during breast cancer progression. A significant clue emanated from our recent study that identified B3GALT6-mediated glycosaminoglycan biosynthesis as a critical dependency required to promote RTC survival in a dormant state (*11*). Clinically, biallelic loss-of-function mutations in *B3GALT6* result in connective tissue disorders with skeletal abnormalities, specifically spondyloepimetaphyseal dysplasia with joint laxity, type 1, and spondylodysplastic Ehlers-Danlos syndrome (*43-46*). Additionally, analogous to the enrichment of sgRNAs targeting *Sox5/6* in the CRISPR-Cas9 screen, we observed an enrichment of sgRNAs targeting *Runx2*, a transcription factor essential for bone development, and for sgRNAs targeting *Evc and Evc2* loss-of-function mutations in which result in the skeletal dwarfism disorder Ellis van Creveld syndrome **(Fig. 1B)**. Indeed, we recently reported the preferential acquisition of *EVC2* missense and putative loss-of-function mutations in metastatic recurrences in breast cancer patients when compared to their matched primary tumors (*47*). Together with our current study, these data support a model in which RTCs adopt developmental programs normally operative in connective tissue and bone as a mechanism of adaptation enabling tumor cell survival as well as maintenance in a dormant state to evade therapy.

Several lines of evidence suggest that the SOX5-driven endochondral ossification program is conserved in different contexts of dormancy and functions to reinforce the dormant state. First, we identified osteochondrogenic differentiation in multiple therapy-associated contexts. These include two independent genetically engineered mouse (GEM) models for HER2-dependent and WNT1-dependent mammary tumors, wherein residual lesions selectively enrich osteochondrogenic gene expression following oncogene inhibition simulating targeted therapy. Critically, we identified similar enrichment of osteochondrogenic gene expression in residual lesions in breast cancer patients following neoadjuvant endocrine therapy or chemotherapy that was independent of breast cancer subtype and therapy modality. Second, we demonstrated osteochondrogenic differentiation and SOX5 activity in contexts where dormancy is induced in tumor cells by exposure to a foreign microenvironment. These include HER2-dependent mammary tumor cells in the lung as well as D2.OR cells in a growth-restrictive 3D extracellular matrix lacking collagen I. Third, we found that enhanced osteochondrogenic gene expression in primary breast cancers in patients, or in residual disease following neoadjuvant therapy, was strongly associated with improved relapse-free survival. Because the vast majority of patient tumors in this dataset were ER+ and experienced metastatic recurrence, these findings suggest that osteochondrogenic differentiation may be broadly recapitulated as an adaptive mechanism independently of breast cancer subtype or the particular context that induced acquisition of a dormant state.

A phenomenon termed osteomimicry has previously been described in breast cancer (*48*) wherein cells acquire osteoblast or osteoclast markers, which enables their metastasis to bone (*49, 50*). In contrast to osteomimicry, which confers pro-tumorigenic functions such as survival, invasion, and proliferation, our data indicate that osteochondrogenic differentiation in RTCs in the mammary gland and the lung promotes a dormant state that is associated with indolent tumor progression. Furthermore, neither SOX5 nor SOX6 have been implicated in osteomimicry. For these reasons, and others, we believe that the acquisition of osteogenic features in dormant MRD is functionally distinct from osteomimicry and part of a broader program that more closely resembles cartilage-dependent bone development.

In summary, our study identifies a role for a SOX5-driven osteochondrogenesis-like pathway in the sustenance of dormant RTCs and provides evidence for its potential as a tool to predicting recurrence risk in breast cancer patients. Future studies incorporating [^18^F]NaF-PET in patients following neoadjuvant therapy could ostensibly be used to prospectively identify patients whose residual lesions adopt osteochondrogenic differentiation, thus suggesting a low likelihood of recurrence. By stratifying patients at risk of recurrence, these pathways can help pinpoint those most likely to benefit from emerging adjuvant MRD-targeting therapies, which has the potential to transform treatment options for millions of breast cancer survivors.

## Acknowledgments

We thank Jianping Wang for histology assistance, Yan Chen for cloning assistance, as well as other members of the Chodosh laboratory for their invaluable input that helped develop this study.

## Funding

National Institutes of Health grant R01CA098371 (LAC)

National Institutes of Health grant R01CA143296 (LAC)

National Institutes of Health grant R01 CA143296 (LAC)

Breast Cancer Research Foundation BCRF-24-026 (LAC)

Susan G. Komen SAC232145 (DAM)

Department of Defense W81XWH-20-1-0013 (AS)

Philanthropy: Rhoda Polly Danziger and Michael Danziger (LAC)

## Author contributions

Conceptualization: AS, DAM, LAC

Methodology: AS, EB, TP, DKS, SA, HT, SDC, MAS, DAM, LAC

Investigation: AS, EB, CJS, TP, GKB, SDC, CAA

Visualization: AS, TP, DKP

Funding acquisition: DAM, LAC

Writing: AS, MAS, DAM, LAC

## Competing interests

LAC has served as an expert consultant to Teva Pharmaceuticals, Eisai, Sanofi, Eli Lilly, Whittaker, Clark and Daniels, Wyeth, Imerys, Colgate, Becton Dickinson, Sterigenics, and the U.S. Department of Justice in litigation. The remaining authors declare no competing interests.

## Data and materials availability

All data are available in the main text or the supplementary materials.

## Materials and Methods

### In vivo experiments

All animal studies received approval from the Institutional Animal Care and Use Committee (IACUC) at the University of Pennsylvania. This study utilized 5–6-week-old female *nu/nu* mice (NCRNU-F, Taconic) as transplantation donors. The mice were ear-tagged and randomly assigned to experimental groups, housed in a barrier facility under 12-hour light/12-hour dark cycles, with ad libitum access to food and water.

Recurrence-free survival assays were carried out following established protocols. Briefly, 1 × 10⁶ primary *MTB/TAN* Her2-dependent tumor cells were transplanted into the #4 inguinal mammary fat pads of female nu/nu mice (NCRNU-F, Taconic). The mice received doxycycline (2 mg/ml) and 5% sucrose through their drinking water. Tumor deinduction was achieved by switching the mice to regular drinking water when the mammary tumors reached the target size (8 × 8 mm for longitudinal PET/CT imaging; 3 × 3 mm for recurrence-free survival assays and the CRISPR-Cas9 screen). Mice were palpated three times weekly, and recurrence-free survival was monitored using Kaplan-Meier analysis, with time-to-recurrence determined by the re-emergence of palpable tumors from non-palpable, dormant lesions. Two hours prior to euthanasia, mice were given an intraperitoneal injection of EdU at a dose of 50 mg/kg.

Experimental metastasis recurrence-free survival was performed by injecting 1 × 10^4^ primary Her2-dependent tumor cells transduced with a vector expressing *Renilla* luciferase and introduced to the lung via tail-vein injection into female *nu/nu* mice (NCRNU-F, Taconic). The mice received doxycycline (2 mg/ml) and 5% sucrose through their drinking water. Her2-dependent tumor growth was monitored weekly using bioluminescence imaging by Firefly luciferase, which is downstream of the Her2 transgene. Mice were switched to regular drinking water to induce tumor deinduction when Firefly luciferase signal exceeded a total flux of 10^9^ photons/s. Following deinduction, tumor burden and time-to-spontaneous regrowth was monitored weekly using bioluminescence for *Renilla* luciferase.

### In vitro assays

Primary Her2-dependent tumor cells were derived from female *MTB/TAN* FVB transgenic mice, as previously described (*10*). The cells were cultured in DMEM (Corning, Cat. #10-017-CV), supplemented with 10% bovine calf serum (Cytiva, Cat. #SH30072.03), 1% Penicillin/Streptomycin (Thermo Fisher Scientific, Cat. #15-140-122), 1% Glutamine (Thermo Fisher Scientific, Cat. #25030081), 2 mg/ml Doxycycline (RPI, Cat. #D43020-250.0), 5 mg/ml Prolactin (NHPP, Cat. #NIDDK-oPRL-21), 5 mg/ml Insulin (GeminiBio, Cat. #700-112P), 10 μg/ml EGF (Millipore, Cat. #E4127), 1 mg/ml Hydrocortisone (Sigma, Cat. #H0396), and 1 mM Progesterone (Sigma, Cat. #P7556).

For in vitro dormancy experiments, cells were plated on Day -3 in the aforementioned medium, transitioned on Day -2 to medium containing 1% bovine calf serum, and further transitioned to medium with 1% bovine calf serum but no doxycycline on Day 0. Plates were harvested at specified dormancy deinduction time points, while medium containing 1% bovine calf serum without doxycycline was replenished weekly for remaining plates. Reinduction plates were harvested at the desired time points after 48-hour treatments with medium containing 1% bovine calf serum and doxycycline, as depicted in the figures.

D2.OR cells, generously provided by Dr. Mikala Egeblad at Cold Spring Harbor Laboratories, originate in *Balb/c* mice. These cells were cultured in DMEM (Corning, Cat. #10-017-CV) with 10% fetal bovine serum (Cytiva, Cat. #SH30910.03) and 1% Penicillin/Streptomycin (Thermo Fisher Scientific, Cat. #15-140-122).

For 3D dormancy studies, D2.OR cells were plated on matrices, composed of either Cultrex 3D Basement Membrane Extract, Reduced Growth Factor (Trevigen, Cat. #3445-005-01) or a 1:1 mixture of basement membrane extract and neutralized type I collagen (Cultrex 3D Culture Matrix Rat Collagen I, Trevigen, Cat. #3447-020-01). For experiments conducted in 96-well plates (Corning, Cat. #353219), 50 μl of matrix was added per well and solidified at 37°C for 1 hour. The cells were resuspended at 20,000 cells/ml in DMEM low glucose, low pyruvate medium (Thermo Fisher Scientific, Cat. #11885092), supplemented with 2% fetal bovine serum (GeminiBio, Cat. #100-106), 2% basement membrane extract, and 1% Penicillin/Streptomycin (Thermo Fisher Scientific, Cat. #15-140-122). A volume of 100 μl per well was then plated on top of the solidified matrices. Plates were harvested at specified time points and medium was refreshed every 4 days for the remaining plates.

Cell viability assays, performed in both 2D and 3D formats, followed the manufacturer’s instructions using the Cell Titer 96 Non-Radioactive Cell Proliferation Assay (Promega, Cat. #G4000) in 96-well plates.

All cells were verified to be free of *Mycoplasma* contamination.

### Plasmids and lentiviral production

LentiV_Cas9_puro (Addgene, Cat. #108100) was utilized to create Cas9-expressing *MTB/TAN*-derived primary tumor cells. For CRISPR-Cas9 studies, sgRNA cloning was performed using LRG2.1 (Addgene, Cat. #108098), LRG (Addgene, Cat. #65656), or custom-cloned LR-Rluc-P2A-GFP vector backbones. Sense and antisense oligos for each sgRNA were phosphorylated, annealed, and ligated into BsmB1-digested vectors. The ligated vectors were transformed into chemically competent Stbl3 bacteria (Thermo Fisher Scientific, Cat. #C737303). Transformed bacterial colonies were selected on Ampicillin plates, and DNA was isolated and sequenced with a U6 primer to confirm sgRNA integration.

Lentivirus production was carried out in HEK293T cells using TransIT-293 transfection reagent (Mirus, Cat. #MIR2700). Packaging plasmids pMD2.G (Addgene, Cat. #12259; 3 μg), psPAX2 (Addgene, Cat. #12260; 6 μg), and 9 μg of the backbone containing the sgRNA/shRNA of interest were introduced. Lentiviral titers were quantified by serial dilution in *MTB/TAN*-derived primary tumor cells, leveraging the fluorophore linked to the vector backbones as a detection marker via Attune NxT flow cytometry (Thermo Fisher).

### PCR and ICE analysis

To validate sgRNA editing efficiency, genomic DNA was isolated from Her2-dependent Cas9 unedited and edited cells one week after transduction with sgRNAs (Qiagen, QIAamp DNA mini kit, Cat. # 51304). PCR amplification was performed (98°C 5min → (98°C 5s, <u>T</u>_m_ 5s, 72°C 15s) ×30 → 72°C 3min → 4°C) using the following primer sequences and annealing temperatures (T_m_):

sg*Rosa*:

F-GCGGGAGAAATGGATATGAA; R-GCACTTGCTCTCCCAAAGTC; T_m_ = 60°C

sg*Sox5*_5:

F- TGAAATGTTCCCACACCCGA; R- AGTCATCCTCTCAGCCTGGT; T_m_ = 64.9°C

sg*Sox5*_3:

F- ATGCTTCTTCCAAGGAGGGC; R- ATCTGCCTTAGGAACTGCGC; T_m_ = 65.4°C

sg*Sox6*_1:

F- CCTGAGCTGTGGTGTGGTAG; R- GAAGATGGGCCAGAGGACAG; T_m_ = 65.1°C

sg*Sox6*_4:

F- CACCTGCTTCTCAGTGCTCA; R- TGCAGCTGAAAGGACTGAGG; T_m_ = 65.1°C

The resulting amplicons were Sanger sequenced and ICE analysis was performed on the sequences to determine editing efficiency using https://ice.synthego.com/#/

### In vivo CRISPR-Cas9 screen

From the genes selectively enriched during dormancy in vitro and categorized under extracellular matrix-associated gene ontology terms, 95 were chosen for CRISPR-Cas9 in vivo screening. The library also included positive control sgRNAs targeting genes known to promote disease progression (pro-proliferative and pro-survival genes), as well as negative control sgRNAs that were non-targeting. For the selected 95 genes, 4–5 sgRNAs were designed for each, resulting in a total of 509 sgRNAs (Table S1). A size-matched library of non-targeting sgRNAs was also created to assist with data analysis.

sgRNAs were designed to target conserved functional domains while minimizing off-target effects, as predicted by the GUIDES tool (http://guides.sanjanalab.org/). Preference was given to sgRNAs with an A/T nucleotide at the 17th position of the sequence to optimize functionally ‘null’ mutations and eliminate the need for subcloning, thus preserving cell heterogeneity. These sgRNA oligo pools were cloned into the LRG expression vector, after which the plasmid pool was amplified, purified, and packaged into lentiviruses. To ensure that MTB/TAN-Cas9 cells received only one sgRNA per cell, the lentiviral library was titrated using the GFP selectable marker and assessed via flow cytometry (Attune NxT, Thermo Fisher), then transduced at an MOI = 0.3. Subsequently, the transduced cells were sorted (MoFlo Astrios, Beckman Coulter Life Sciences), expanded, and prepared for in vivo transplantation. All procedures ensured that each sgRNA was represented in >1500 cells/sgRNA to maintain adequate coverage and enhance the robustness of downstream analyses.

Mouse primary tumors and residual lesions were harvested, microdissected under a stereoscope, and homogenized for genomic DNA extraction using the Quick-DNA Midiprep Plus kit (Zymo Research, Cat. #D4075). sgRNA inserts were PCR-amplified using the Phusion Flash High Fidelity PCR Master Mix (Thermo Fisher, Cat. #F548), followed by the addition of barcodes and adapters. The libraries were pooled, supplemented with 5% PhiX (Thermo Fisher, Cat. #FC-110-3001), and sequenced in parallel using the MiSeq reagent kit v3 (150 cycles) (Illumina, Cat. #MS-102-3001) on the MiSeq Instrument (Illumina).

Fastq files containing sequencing reads were demultiplexed using in-house R script, followed by sgRNA counting using modified version of a published Python script (*51*). Percent abundance was calculated for each sgRNA relative to total read count per sample. Read counts from technical replicates were combined. Diversity in read distribution across sgRNAs in a sample was visualized by plotting the cumulative percentage of reads against descending ranks of sgRNAs by read counts for comparison across different samples and time points. Potential clonal or emerging clonal genes were identified by looking for sgRNAs satisfying at least one of the following: 1) percent abundance in a sample is >5%, 2) abundance is more than k x interquartile range above the upper quartile (k is a multiplier described below), or 3) abundance in a sample is significantly above the levels in primary tumor samples as determined using edgeR (*52*) with a fold change cutoff of 2 and an FDR cutoff of 0.1. For the second criterion, the multiplier k was defined as the minimum value resulting in fewer than 1 outlier sgRNA per primary tumor sample in the data set.

### PET imaging and analysis

Mice were injected with approximately 250 uCi of [^18^F]NaF. At 1 h post injection, the mice were placed into an induction chamber with 3% isoflurane. After being anesthetized, the mice were placed on the bed of the Molecubes β-Cube PET scanner (Molecubes NV, Gent, Belgium) and imaged for 10 minutes with continuous isoflurane anesthesia. A general purpose CT scan on the Molecubes X-Cube (Molecubes NV, Gent, Belgium) followed for data correction and anatomical reference. After imaging, the mice were recovered. At endpoint, mice were euthanized and tumors were dissected from the animals and imaged. For the dissected tumors, the samples were placed into plastic wells filled with Tissue-Tek O. C. T. Compund (Sakura, Cat. #4583). The wells were then placed onto a bed and scanned with the PET scanner for 10 minutes followed by a CT scan.

PET image analysis was performed using Amide (A Medical Image Data Examiner) software. [^18^F]NaF uptake was calculated as follows. Whole animal ROIs were drawn as boxes with dimensions x = 35, y = 120, z = 25, whereas mammary gland ROIs were generated as an elliptical cyclinder with dimensions x = 2, y = 2, z = 2 and manually adjusted to encompass the lesion as detected on the CT image. Mean values generated by Amide in kBq/cc were multiplied by the total volume of ROIs (in cc) to generate the Sum values in kBq and to generate % injected dose in the mammary gland/uptake in the whole animal in each bilateral site/mouse.

### Histology

For in vitro immunofluorescence and labeling studies, MTB/TAN-derived primary tumor cells were cultured on glass coverslips (Bellco Glass Inc., Cat. #1943-010015A) and treated with 5 μM EdU for 2 hours before harvest. Coverslips were fixed with 4% paraformaldehyde for 15 minutes at room temperature, permeabilized with PBS containing 0.5% Triton X-100 for 20 minutes at room temperature, and washed with 3% bovine serum albumin in 1× PBS before proceeding with the immunofluorescence protocol.

For in vivo immunofluorescence and labeling studies, inguinal #4 mammary fat pads containing primary tumors, residual lesions, or recurrent tumors were dissected and fixed in 4% paraformaldehyde overnight at 4°C. Samples were then thoroughly washed, dehydrated, embedded in paraffin, and sectioned at 5 μm. Slides were prepared by serial deparaffinization and rehydration, followed by PBS washes and antigen retrieval using R-Buffer A (Electron Microscopy Sciences, Cat. #62706-10) or R-Buffer B (Electron Microscopy Sciences, Cat. #62706-11) in a retriever (Aptum, Cat. #RR2100-EU).

EdU labeling studies were conducted on coverslips (Thermo Fisher Scientific, Cat. #C10640) or tissues (Sigma Aldrich, Cat. #BCK647-IV-IM-S) processed as per manufacturer’s instructions.

Immunofluorescence samples were blocked with 5% goat serum in 1× PBS with mouse-on-mouse block (Vector Laboratories, Cat. #BMK-2202). After three washes in 1× PBS, samples were incubated overnight at 4°C with primary antibodies or corresponding isotype controls diluted in 5% goat serum in 1× PBS with mouse-on-mouse diluent (Vector Laboratories, Cat. #BMK-2202). The primary antibodies used were: Rat anti-Ki67 (Thermo Fisher Scientific, Cat. #14-5698; 1:100), Rabbit anti-GFP (Cell Signaling, Cat. #2956; 1:200), and Mouse anti-GFP (Living Colors, Cat. #632381; 1:250). Isotype controls included: Rat IgG (Thermo Fisher Scientific, Cat. #02-9602), Rabbit IgG (Thermo Fisher Scientific, Cat. #02-6102), and Mouse IgG2a kappa (Thermo Fisher Scientific, Cat. #14-4724-82).

Following primary antibody incubation, samples were washed three times in 1× PBS and incubated for 1 hour at 37°C with secondary antibodies diluted as follows: Goat anti-mouse IgG2a Alexa488 (Thermo Fisher Scientific, Cat. #A21131; 1:1000), Goat anti-rabbit IgG Alexa488 (Thermo Fisher Scientific, Cat. #A11034; 1:1000), Goat anti-rat IgG Alexa568 (Thermo Fisher Scientific, Cat. #A11077; 1:1000).

For immunofluorescence in the lung, we used the Tyramide SuperBoost Signal Amplification (Thermo Fisher Scientific, Cat. #B40943) protocol with an added hydrogen peroxide quenching step, which removes endogenous peroxidase to increase the efficiency of the amplification.The reaction with HRP (horse radish peroxidase) and hydrogen peroxide drives the activation of a tyramide substrate bound to a fluorophore. The active tyramide substrate binds tyrosine residues around the target protein, amplifying its signal. The reaction is stopped with a stop solution at 5 minutes precisely to maximize signal without increasing the background noise.

Post-secondary antibody incubation, samples were washed three times in 1× PBS and counterstained with 0.5 μg/ml Hoechst 33258 for 10 minutes at room temperature. Samples were mounted using ProLong Gold (Thermo Fisher Scientific, Cat. #P36934) and imaged using a DM 5000B Automated Upright Microscope equipped with a DFC350 FX monochrome digital camera (Leica Microsystems).

### Computational data analysis

CollecTRI (*14*) is a collection of transcription factor regulons (transcription factors and their target genes) with a directionality (mode) of regulation. This can be +1 or -1 depending on whether the target gene is activated or repressed. The CollecTRI regulons include 43175 signed TF-gene interactions from 1186 TFs for human data and 38970 interactions from 1073 TFs for mouse data. Computational algorithms to compute TF activity are provided by the same authors in an R software package called decoupleR (*53*). In our work, we have utilized TF signatures from the CollecTRI collection and plotted standard z-scores.

From our in-house in-vitro dormancy data, we have developed a signature from genes differentially expressed between day 0 and day 28 at an absolute fold change cutoff of 2 and FDR < 0.1 (*11*). Additionally, we have developed a signature from our in-house in-vivo data of residual tumor cells from the Her2 mouse model (*12*). This signature consists of genes differentially expressed between primary tumor cells and residual tumor cells at the same cutoffs above. The intersection of these two signatures constitutes the dormancy signature.

To determine lineage enrichment in the dormancy signature, we mined the mouse organogenesis cell atlas (MOCA), a comprehensive single cell RNAseq experiment of mouse organogenesis by profiling around 2 million cells derived from 61 embryos (*21*). They have identified hundreds of cell types and 56 developmental trajectories. From this resource, the data we have used (cell_annotate.csv, gene_count_cleaned.RDS) is downloaded from the link - (https://oncoscape.v3.sttrcancer.org/atlas.gs.washington.edu.mouse.rna/downloads). To compute the signature scores, the huge gene expression matrix (around 26k genes and 1.4M cells) was reduced to a smaller gene expression matrix and it comprised of 1334 groups of 1000 cells of each cell type. The signature scores are computed per group (of 1000 cells) as z-scores over the signature genes.

Signatures for chondrogenesis and osteogenesis were generated using gene expression data from human fibroblasts grown in osteogenic or chondrogenic medium for 28 days(*22*). The chondrogenesis signature contains 57 genes which were up-regulated in fibroblasts induced for chondrogenic differentiation compared to control fibroblasts (t-test p<0.05, fold change>2). The osteogenesis signature contains 122 genes which were up-regulated in fibroblasts induced for osteogenic differentiation compared to control fibroblasts (t-test p<0.05, fold change>2). To calculate signature scores in human breast cancer data sets (describe below), the log-scale expression levels of each signature gene was z-transformed across all samples within a data set. The signature score for each signature in each sample was then defined as the mean z-score from all signature genes.

Publicly available high-throughput gene expression data for 4,314 patients from 16 human primary breast cancer data sets (*54-69*) along with the corresponding clinical annotations were downloaded from NCBI GEO or original authors’ websites. Within each data set, the effect size (hazard ratio) of the association between signature scores and 5-year relapse-free survival was estimated using Cox proportional hazards regression. The effect size estimates were combined across data sets by meta-analysis using the inverse-variance weighting method(*70*). Overall difference in signature scores across the prognostic subtype groups in the METABRIC data sets was assessed using ANOVA and plotted using relative signature scores for which the baseline was defined as the minimum per-group mean score.

Public gene expression data with matching tumor samples from breast cancer patients before and after neoadjuvant chemotherapy from 10 data sets (*28-30, 32-36, 38, 71, 72*) were downloaded and processed. These represent 395 patients treated with anti-estrogen therapies across 5 studies and 106 patients treated with chemotherapy across 5 studies. Signature scores were calculated in each data set as described above. Mean difference (effect size) in signature scores between pre-treatment and each post-treatment time point was calculated and compared using paired t-test among matched samples. To determine if changes in signatures scores by treatment is associated with breast cancer relapse, we obtained gene expression and clinical data of 21 patients through collaboration with the I-SPY consortium and tested the association using Kaplan-Meier plotter(*73*).

### Quantification and statistical analysis

Primary and recurrent tumor volume data from animal models were converted to tumor size estimates using tumor size = (volume × 6 ÷ π)^1/3^ (i.e. the diameter of a spherical object of equivalent volume). Tumor growth rate was defined as change in tumor size as a linear function of time and was estimated using linear regression between tumor size and time. Tumor growth rates between different experimental groups were compared using Wilcoxon rank-sum test.

For comparisons across multiple groups, ANOVA followed by Sidak’s multiple comparisons test was applied. Kaplan-Meier analysis was conducted for survival curves, with p values calculated using the Gehan-Breslow-Wilcoxon test and hazard ratios calculated using the logrank method.

## References

1. F. Bray et al., Global cancer statistics 2022: GLOBOCAN estimates of incidence and mortality worldwide for 36 cancers in 185 countries. CA Cancer J Clin 74, 229–263 (2024).

2. R. A. Willis, The spread of tumours in the human body. (J. & A. Churchill, London, 1934), pp. x, 540 p.

3. H. Pan et al., 20-Year Risks of Breast-Cancer Recurrence after Stopping Endocrine Therapy at 5 Years. N Engl J Med 377, 1836–1846 (2017).

4. R. N. Pedersen et al., The Incidence of Breast Cancer Recurrence 10-32 Years after Primary Diagnosis. J Natl Cancer Inst, (2021).

5. L. J. Bayne et al., Identifying breast cancer survivors with dormant disseminated tumor cells: The PENN-SURMOUNT screening study [abstract]. Cancer Res 81, (2021).

6. L. J. Bayne et al., Detection and targeting of minimal residual disease in breast cancer to reduce recurrence: The PENN-SURMOUNT and CLEVER trials [abstract]. Cancer Res 78, (2018).

7. E. Dalla, A. Sreekumar, J. A. Aguirre-Ghiso, L. A. Chodosh, Dormancy in Breast Cancer. Cold Spring Harb Perspect Med, (2023).

8. E. J. Gunther et al., A novel doxycycline-inducible system for the transgenic analysis of mammary gland biology. FASEB J 16, 283–292 (2002).

9. S. E. Moody et al., Conditional activation of Neu in the mammary epithelium of transgenic mice results in reversible pulmonary metastasis. Cancer Cell 2, 451–461 (2002).

10. S. E. Moody et al., The transcriptional repressor Snail promotes mammary tumor recurrence. Cancer Cell 8, 197–209 (2005).

11. A. Sreekumar et al., B3GALT6 promotes dormant breast cancer cell survival and recurrence by enabling heparan sulfate-mediated FGF signaling. Cancer Cell 42, 52–69 e57 (2024).

12. J. R. Ruth et al., Cellular dormancy in minimal residual disease following targeted therapy. Breast Cancer Res 23, 63 (2021).

13. V. Lefebvre, The SoxD transcription factors--Sox5, Sox6, and Sox13--are key cell fate modulators. Int J Biochem Cell Biol 42, 429-432 (2010).

14. S. Muller-Dott et al., Expanding the coverage of regulons from high-confidence prior knowledge for accurate estimation of transcription factor activities. Nucleic Acids Res 51, 10934–10949 (2023).

15. L. Gerratana et al., Pattern of metastasis and outcome in patients with breast cancer. Clin Exp Metastasis 32, 125–133 (2015).

16. C. Prunier et al., Breast cancer dormancy is associated with a 4NG1 state and not senescence. NPJ Breast Cancer 7, 140 (2021).

17. A. Zawerton et al., Widening of the genetic and clinical spectrum of Lamb-Shaffer syndrome, a neurodevelopmental disorder due to SOX5 haploinsufficiency. Genet Med 22, 524–537 (2020).

18. V. Lefebvre, Roles and regulation of SOX transcription factors in skeletogenesis. Curr Top Dev Biol 133, 171–193 (2019).

19. V. Lefebvre, R. R. Behringer, B. de Crombrugghe, L-Sox5, Sox6 and Sox9 control essential steps of the chondrocyte differentiation pathway. Osteoarthritis Cartilage 9 Suppl A, S69-75 (2001).

20. P. Smits et al., The transcription factors L-Sox5 and Sox6 are essential for cartilage formation. Dev Cell 1, 277–290 (2001).

21. J. Cao et al., The single-cell transcriptional landscape of mammalian organogenesis. Nature 566, 496–502 (2019).

22. J. Rakar, S. Lonnqvist, P. Sommar, J. Junker, G. Kratz, Interpreted gene expression of human dermal fibroblasts after adipo-, chondro- and osteogenic phenotype shifts. Differentiation 84, 305–313 (2012).

23. E. J. Gunther et al., Impact of p53 loss on reversal and recurrence of conditional Wnt-induced tumorigenesis. Genes Dev 17, 488–501 (2003).

24. M. Blau, W. Nagler, M. A. Bender, Fluorine-18: a new isotope for bone scanning. J Nucl Med 3, 332–334 (1962).

25. I. Fogelman, G. Cook, O. Israel, H. Van der Wall, Positron emission tomography and bone metastases. Semin Nucl Med 35, 135–142 (2005).

26. M. Araz, G. Aras, O. N. Kucuk, The role of 18F-NaF PET/CT in metastatic bone disease. J Bone Oncol 4, 92–97 (2015).

27. O. M. Rueda et al., Dynamics of breast-cancer relapse reveal late-recurring ER-positive genomic subgroups. Nature 567, 399–404 (2019).

28. E. Stickeler et al., Basal-like molecular subtype and HER4 up-regulation and response to neoadjuvant chemotherapy in breast cancer. Oncol Rep 26, 1037–1045 (2011).

29. A. K. Turnbull et al., Accurate Prediction and Validation of Response to Endocrine Therapy in Breast Cancer. J Clin Oncol 33, 2270–2278 (2015).

30. L. M. Arthur et al., Molecular changes in lobular breast cancers in response to endocrine therapy. Cancer Res 74, 5371–5376 (2014).

31. W. R. Miller et al., Changes in breast cancer transcriptional profiles after treatment with the aromatase inhibitor, letrozole. Pharmacogenet Genomics 17, 813–826 (2007).

32. L. A. Korde et al., Gene expression pathway analysis to predict response to neoadjuvant docetaxel and capecitabine for breast cancer. Breast Cancer Res Treat 119, 685–699 (2010).

33. T. Gruosso et al., Chronic oxidative stress promotes H2AX protein degradation and enhances chemosensitivity in breast cancer patients. EMBO Mol Med 8, 527–549 (2016).

34. A. M. Gonzalez-Angulo et al., Gene expression, molecular class changes, and pathway analysis after neoadjuvant systemic therapy for breast cancer. Clin Cancer Res 18, 1109–1119 (2012).

35. A. K. Dunbier et al., Molecular profiling of aromatase inhibitor-treated postmenopausal breast tumors identifies immune-related correlates of resistance. Clin Cancer Res 19, 2775–2786 (2013).

36. C. Selli et al., Molecular changes during extended neoadjuvant letrozole treatment of breast cancer: distinguishing acquired resistance from dormant tumours. Breast Cancer Res 21, 2 (2019).

37. K. Krug et al., Proteogenomic Landscape of Breast Cancer Tumorigenesis and Targeted Therapy. Cell 183, 1436–1456 e1431 (2020).

38. R. J. Bownes et al., On-treatment biomarkers can improve prediction of response to neoadjuvant chemotherapy in breast cancer. Breast Cancer Res 21, 73 (2019).

39. L. J. Esserman et al., Chemotherapy response and recurrence-free survival in neoadjuvant breast cancer depends on biomarker profiles: results from the I-SPY 1 TRIAL (CALGB 150007/150012; ACRIN 6657). Breast Cancer Res Treat 132, 1049–1062 (2012).

40. T. Saphner, D. C. Tormey, R. Gray, Annual hazard rates of recurrence for breast cancer after primary therapy. J Clin Oncol 14, 2738–2746 (1996).

41. M. Colleoni et al., Annual Hazard Rates of Recurrence for Breast Cancer During 24 Years of Follow-Up: Results From the International Breast Cancer Study Group Trials I to V. J Clin Oncol 34, 927–935 (2016).

42. D. W. Cescon et al., Therapeutic Targeting of Minimal Residual Disease to Prevent Late Recurrence in Hormone-Receptor Positive Breast Cancer: Challenges and New Approaches. Front Oncol 11, 667397 (2021).

43. A. Ben-Mahmoud et al., A B3GALT6 variant in patient originally described as Al-Gazali syndrome and implicating the endoplasmic reticulum quality control in the mechanism of some beta3GalT6-pathy mutations. Clin Genet 93, 1148–1158 (2018).

44. S. Delbaere et al., b3galt6 Knock-Out Zebrafish Recapitulate beta3GalT6-Deficiency Disorders in Human and Reveal a Trisaccharide Proteoglycan Linkage Region. Front Cell Dev Biol 8, 597857 (2020).

45. F. Malfait et al., Defective initiation of glycosaminoglycan synthesis due to B3GALT6 mutations causes a pleiotropic Ehlers-Danlos-syndrome-like connective tissue disorder. Am J Hum Genet 92, 935–945 (2013).

46. M. Nakajima et al., Mutations in B3GALT6, which encodes a glycosaminoglycan linker region enzyme, cause a spectrum of skeletal and connective tissue disorders. Am J Hum Genet 92, 927–934 (2013).

47. M. R. Paul et al., Genomic landscape of metastatic breast cancer identifies preferentially dysregulated pathways and targets. J Clin Invest 130, 4252–4265 (2020).

48. O. Awolaran, S. A. Brooks, V. Lavender, Breast cancer osteomimicry and its role in bone specific metastasis; an integrative, systematic review of preclinical evidence. Breast 30, 156–171 (2016).

49. C. C. Tan et al., Breast cancer cells obtain an osteomimetic feature via epithelial-mesenchymal transition that have undergone BMP2/RUNX2 signaling pathway induction. Oncotarget 7, 79688–79705 (2016).

50. S. Wang et al., FOXF2 reprograms breast cancer cells into bone metastasis seeds. Nat Commun 10, 2707 (2019).

51. J. Joung et al., Genome-scale CRISPR-Cas9 knockout and transcriptional activation screening. Nat Protoc 12, 828–863 (2017).

52. M. D. Robinson, D. J. McCarthy, G. K. Smyth, edgeR: a Bioconductor package for differential expression analysis of digital gene expression data. Bioinformatics 26, 139–140 (2010).

53. I. M. P. Badia, et al., decoupleR: ensemble of computational methods to infer biological activities from omics data. Bioinform Adv 2, vbac016 (2022).

54. H. Y. Chang et al., Robustness, scalability, and integration of a wound-response gene expression signature in predicting breast cancer survival. Proc Natl Acad Sci U S A 102, 3738–3743 (2005).

55. M. Chanrion et al., A gene expression signature that can predict the recurrence of tamoxifen-treated primary breast cancer. Clin Cancer Res 14, 1744–1752 (2008).

56. K. Chin et al., Genomic and transcriptional aberrations linked to breast cancer pathophysiologies. Cancer Cell 10, 529–541 (2006).

57. C. Curtis et al., The genomic and transcriptomic architecture of 2,000 breast tumours reveals novel subgroups. Nature 486, 346–352 (2012).

58. C. Desmedt et al., Strong time dependence of the 76-gene prognostic signature for node-negative breast cancer patients in the TRANSBIG multicenter independent validation series. Clin Cancer Res 13, 3207–3214 (2007).

59. K. R. Hess et al., Pharmacogenomic predictor of sensitivity to preoperative chemotherapy with paclitaxel and fluorouracil, doxorubicin, and cyclophosphamide in breast cancer. J Clin Oncol 24, 4236–4244 (2006).

60. A. V. Ivshina et al., Genetic reclassification of histologic grade delineates new clinical subtypes of breast cancer. Cancer research 66, 10292–10301 (2006).

61. X. J. Ma et al., A two-gene expression ratio predicts clinical outcome in breast cancer patients treated with tamoxifen. Cancer Cell 5, 607–616 (2004).

62. A. J. Minn et al., Lung metastasis genes couple breast tumor size and metastatic spread. Proc Natl Acad Sci U S A 104, 6740–6745 (2007).

63. D. S. Oh et al., Estrogen-regulated genes predict survival in hormone receptor-positive breast cancers. J Clin Oncol 24, 1656–1664 (2006).

64. Y. Pawitan et al., Gene expression profiling spares early breast cancer patients from adjuvant therapy: derived and validated in two population-based cohorts. Breast Cancer Res 7, R953–964 (2005).

65. R. Sabatier et al., A gene expression signature identifies two prognostic subgroups of basal breast cancer. Breast cancer research and treatment 126, 407–420 (2011).

66. M. Schmidt et al., The humoral immune system has a key prognostic impact in node-negative breast cancer. Cancer research 68, 5405–5413 (2008).

67. C. Sotiriou et al., Gene expression profiling in breast cancer: understanding the molecular basis of histologic grade to improve prognosis. Journal of the National Cancer Institute 98, 262–272 (2006).

68. Y. Wang et al., Gene-expression profiles to predict distant metastasis of lymph-node-negative primary breast cancer. Lancet 365, 671–679 (2005).

69. L. J. Esserman et al., Chemotherapy response and recurrence-free survival in neoadjuvant breast cancer depends on biomarker profiles: results from the I-SPY 1 TRIAL (CALGB 150007/150012; ACRIN 6657). Breast cancer research and treatment 132, 1049-1062 (2012).

70. A. Ramasamy, A. Mondry, C. C. Holmes, D. G. Altman, Key issues in conducting a meta-analysis of gene expression microarray datasets. PLoS Med 5, e184 (2008).

71. M. Anurag et al., Immune Checkpoint Profiles in Luminal B Breast Cancer (Alliance). Journal of the National Cancer Institute 112, 737–746 (2020).

72. W. R. Miller, A. Larionov, Changes in expression of oestrogen regulated and proliferation genes with neoadjuvant treatment highlight heterogeneity of clinical resistance to the aromatase inhibitor, letrozole. Breast Cancer Res 12, R52 (2010).

73. B. Gyorffy, Integrated analysis of public datasets for the discovery and validation of survival-associated genes in solid tumors. Innovation (Camb) 5, 100625 (2024).

