## Supplementary figures for "Residual Breast Cancer Cells Co-opt SOX5-driven Endochondral Ossification to Maintain Dormancy"

### Figure S1

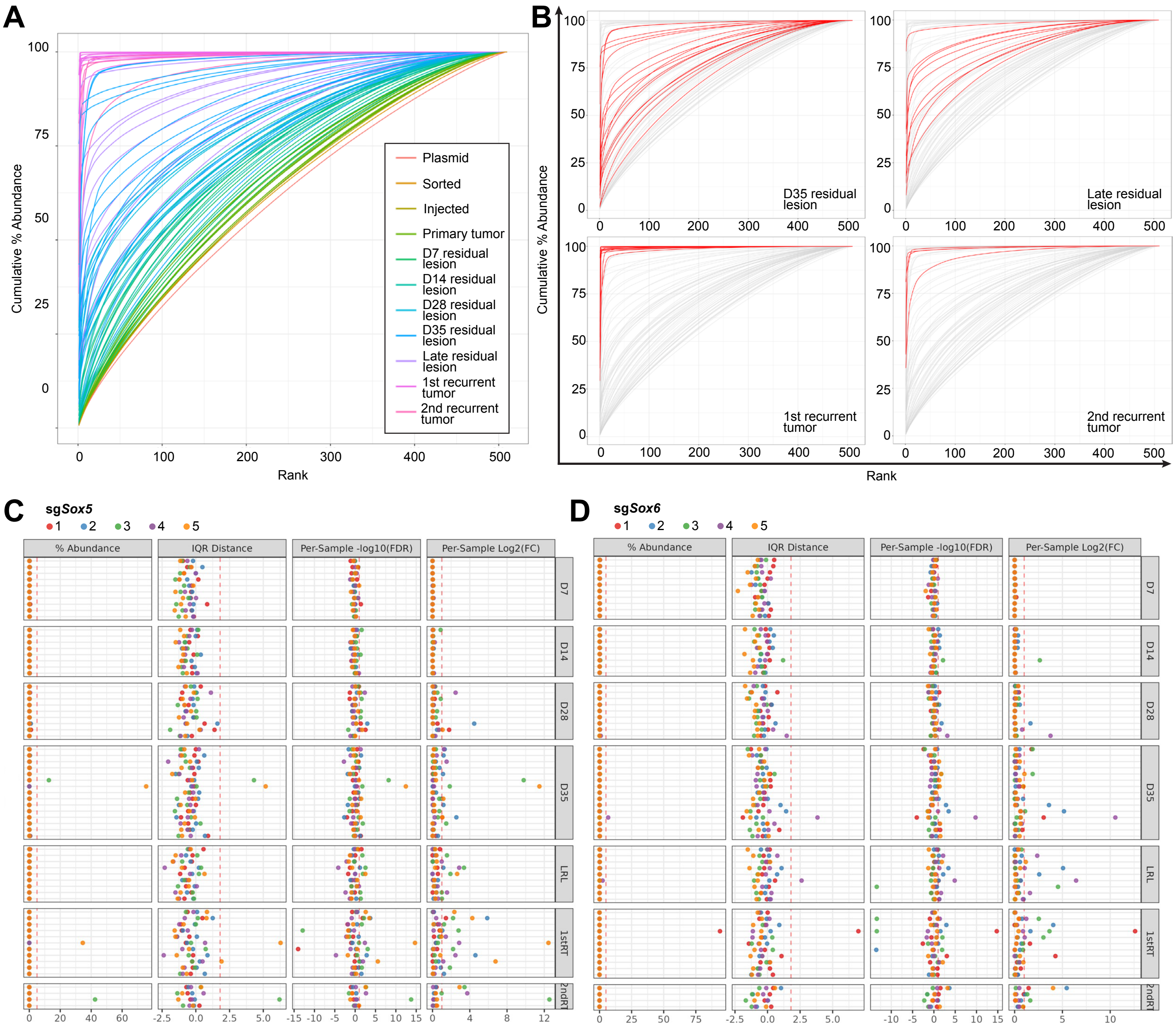


**Fig S1: A.** Cumulative abundance of 509 sgRNAs at sequential time points during disease progression colored by time points. **B.** Cumulative abundance of 509 sgRNAs at late residual lesion time points and recurrent tumors. **C.** Statistical methods for identifying clonal enrichment of sgRNAs targeting *Sox5* and **D.** *Sox6*. Cutoffs indicated by red dotted lines - high abundance > 5%, interquartile (IQR) distance > 1.8, false discovery rate (FDR) < 0.1, fold change (FC) >2.

### Figure S2

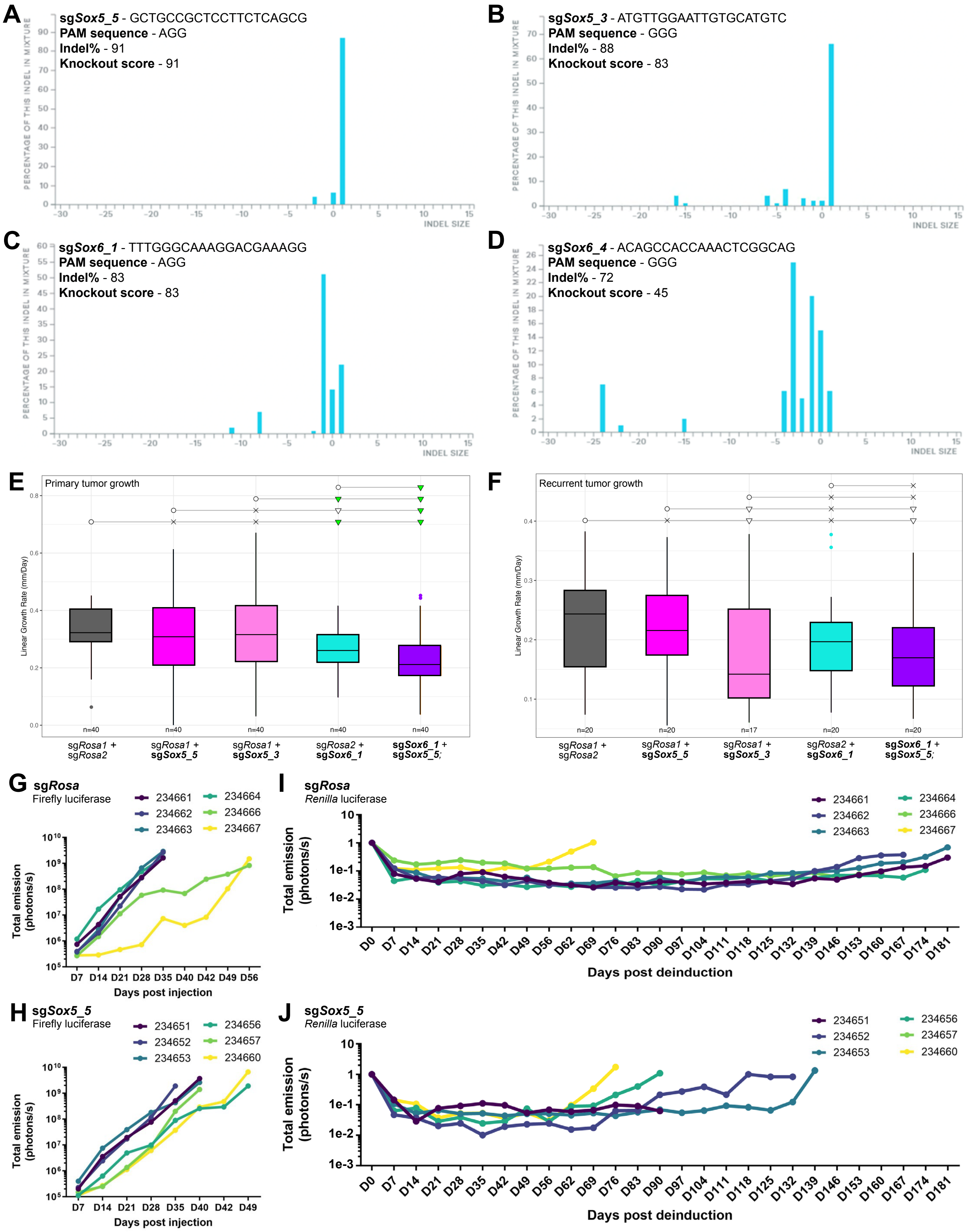

A
sgSox5_5 - GCTGCCGCTCCTTCTCAGCG
PAM sequence - AGG
Indel% - 91
Knockout score - 91
PERCENTAGE OF THIS INDEL IN MIXTURE
INDEL SIZE
B
sgSox5_3 - ATGTTGGAATTGTGCATGTC
PAM sequence - GGG
Indel% - 88
Knockout score - 83
C
sgSox6_1 - TTTGGGCAAAGGACGAAAGG
PAM sequence - AGG
Indel% - 83
Knockout score - 83
D
sgSox6_4 - ACAGCCACCAAACTCGGCAG
PAM sequence - GGG
Indel% - 72
Knockout score - 45
E
Primary tumor growth
Linear Growth Rate (mm/Day)
n=40
sgRosa1 + sgRosa2
sgRosa1 + sgSox5_5
sgRosa1 + sgSox5_3
sgRosa2 + sgSox6_1
sgSox6_1 + sgSox5_5;
F
Recurrent tumor growth
Linear Growth Rate (mm/Day)
n=20
n=17
G
sgRosa
Firefly luciferase
234661
234662
234663
234664
234666
234667
Total emission (photons/s)
Days post injection
I
sgRosa
Renilla luciferase
Total emission (photons/s)
Days post deinduction
H
sgSox5_5
Firefly luciferase
234651
234652
234653
234656
234657
234660
Total emission (photons/s)
Days post injection
J
sgSox5_5
Renilla luciferase
Total emission (photons/s)
Days post deinduction

### Figure S2

**Fig S2: A.** Inference of CRISPR edits (ICE) analysis for sg*Sox5*_5, **B.** sg*Sox5*_3, **C.** sg*Sox6*_1, and **D.** sg*Sox6*_4 displaying the distribution of indels in the population as a function of indel size. **E.** Change in primary tumor or **F.** recurrent tumor size as a linear function of time (mm/day), estimated using linear regression between tumor size and days of growth. Circle = baseline, x = non-significant, empty triangles = trending significance $0.05 < p < 0.1$, filled triangles = $p < 0.05$. **G.** Firefly luciferase total emission (photons/s) during tumor growth till doxycycline withdrawal in mice injected with sg*Rosa* or **H.** sg*Sox5*_5 cells. **I.** Normalized *Renilla* luciferase total emission (photons/s) following doxycycline withdrawal in mice injected with sg*Rosa* or **J.** sg*Sox5*_5 cells.

### Figure S3

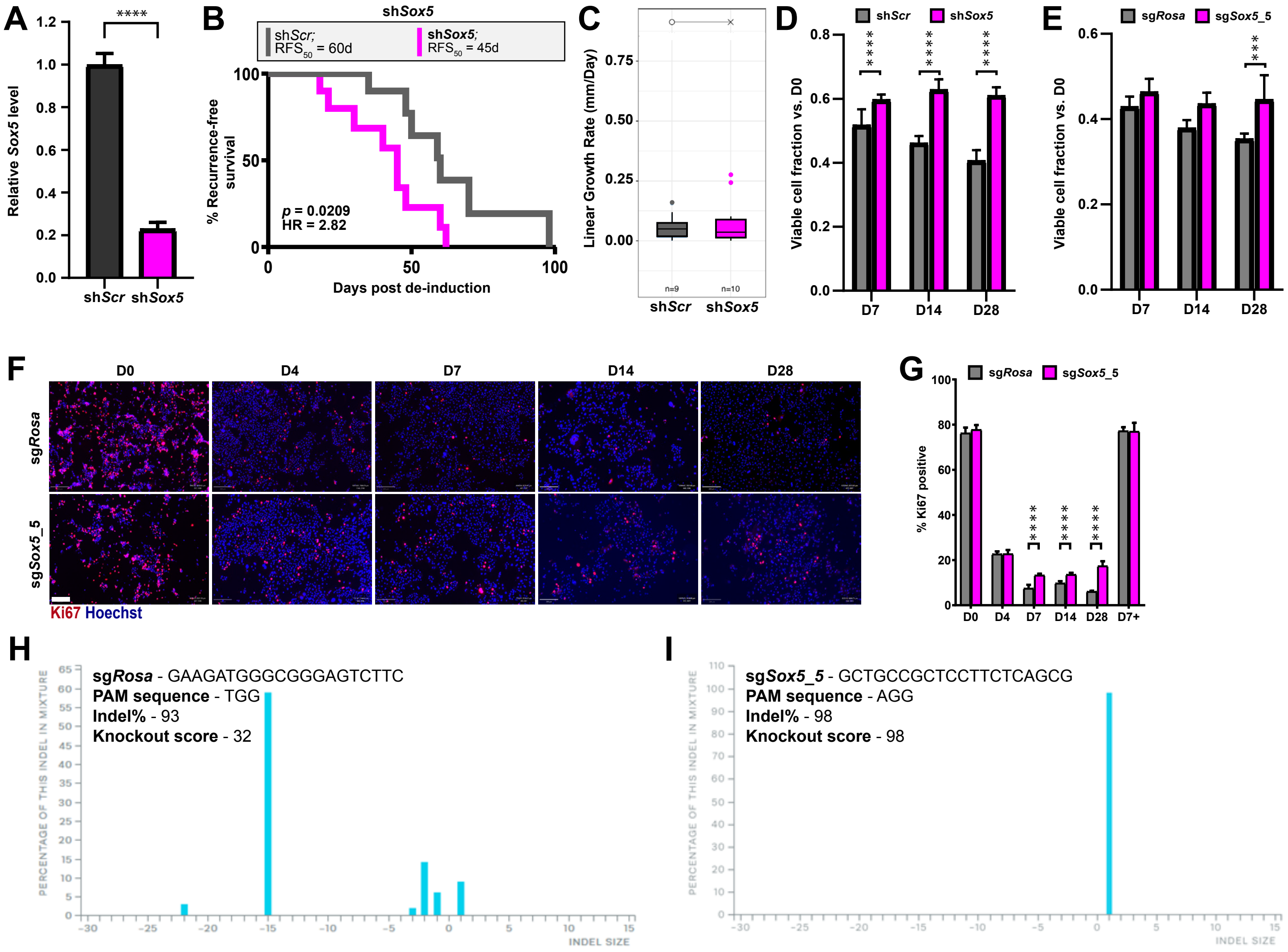


**Fig S3: A.** qRT-PCR for *Sox5* transcripts in sh*Scr* or sh*Sox5* *MTB/TAN* cells. ****$p < 0.0001$. **B.** Kaplan-Meier analysis of recurrence-free survival for sh*Scr* (gray) and sh*Sox5* (fuchsia) groups. **C.** Change in recurrent tumor size as a linear function of time (mm/day), estimated using linear regression between tumor size and days of growth. Circle = baseline, x = non-significant. **C.** Change in recurrent tumor size as a linear function of time (mm/day), estimated using linear regression between tumor size and days of growth. Circle = baseline, x = non-significant. **D.** Viable RTC counts relative to D0 in sh*Scr* (gray) and sh*Sox5* (fuchsia) *MTB/TAN* cells and **E.** sg*Rosa* (gray) and sg*Sox5*_5 (fuchsia) *MTB/TAN*-Cas9 cells. Quantification is represented as mean ± SD. n = 6 replicates/group. ***$p < 0.001$, ****$p < 0.0001$. **F.** Immunofluorescence and **G.** quantification for Ki67 (red) in tumor cells (green) in g*Rosa* (gray) and sg*Sox5*_5 (fuchsia) *MTB/TAN*-Cas9 cells in vitro. Quantification is represented as mean ± SD. n = 3 replicates/group. ****$p < 0.0001$. Scale bar = 200µm. **H.** Inference of CRISPR edits (ICE) analysis for sg*Rosa* and **I.** sg*Sox5*_5 displaying the distribution of indels in the population as a function of indel size.

#### Figure S4

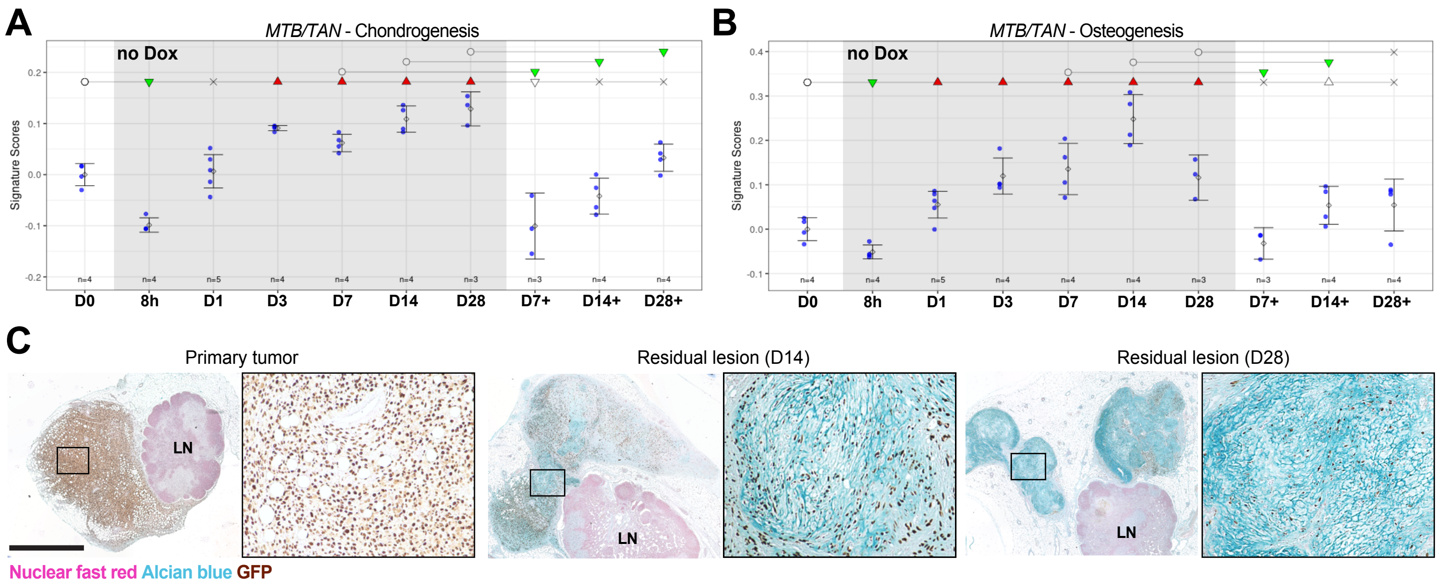


**Fig S4: A.** Application of the chondrogenesis and **B.** osteogenesis signatures on in vitro dormancy time points using the *MTB/TAN* cells. Circle = baseline, x = non-significant, empty triangles = trending significance $0.05 < p < 0.1$, filled triangles = $p < 0.05$. **C.** Near-adjacent sections relative to those in Fig. 4G displaying GFP+ tumor cells (brown) relative to Alcian blue staining in primary tumor and residual lesion sections. LN = lymph node. Scale bar = 2mm.

### Figure S5

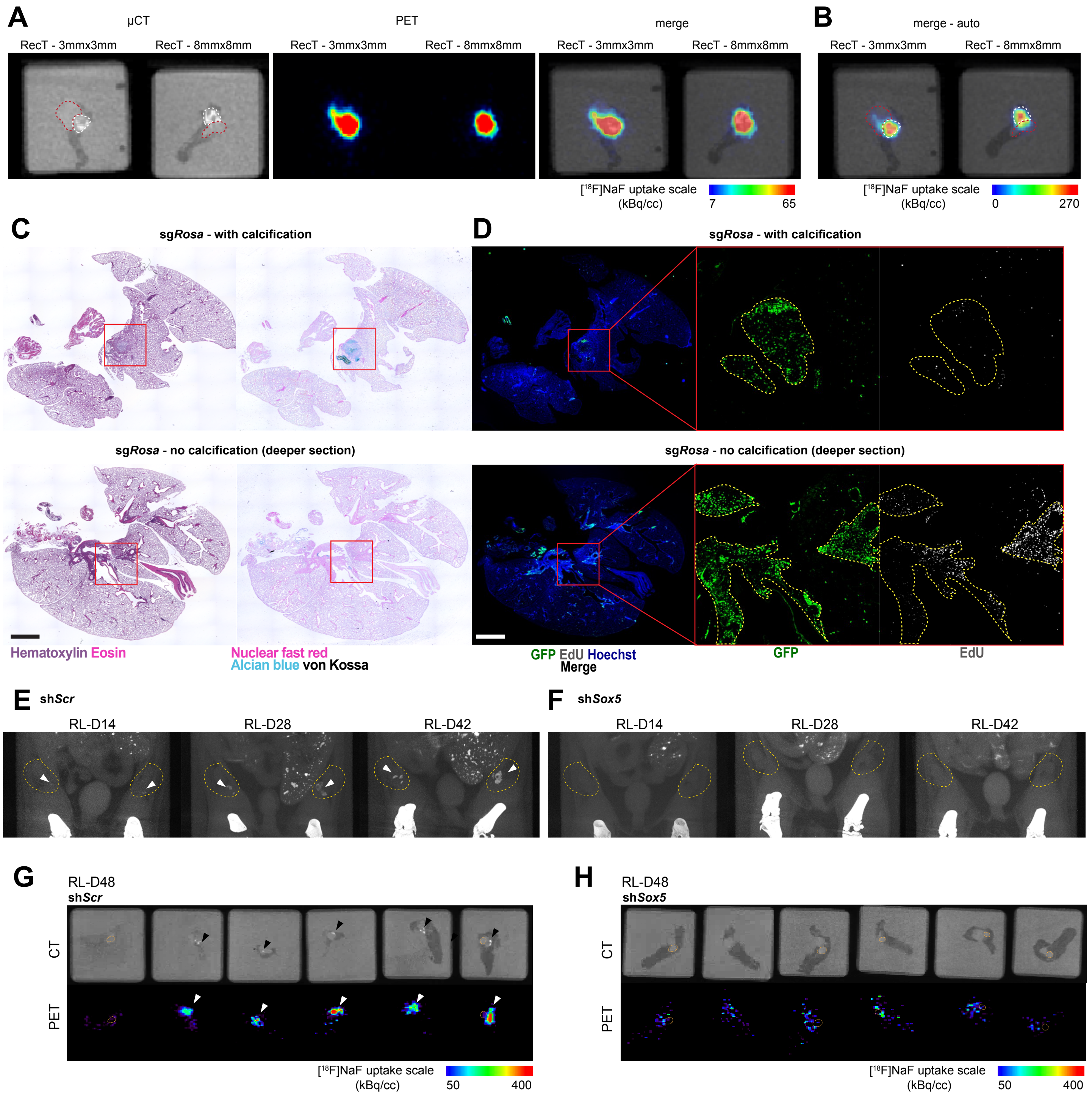


**Fig S5: A.** µCT and PET 75 minutes after injection of [$^{18}$F]NaF. Calcified areas (white dotted line) and non-calcified recurrent outgrowth (red dotted line) indicated on µCT images with comparable scaling to Fig. 5C or **B.** auto scaling. **C.** Serial sections of a recurrent tumors in the lung stained with Hematoxylin and Eosin (panel 1) as well as Alcian blue (blue) and von Kossa (black) to identify cartilage- and bone-like areas, respectively (panel 2). Red boxes indicate regions of interest. **D.** Immunofluorescence for GFP+ tumor cells (green), 24h EdU uptake (gray) in regions of interest (red box), with magnified views. Yellow dotted lines indicate regions containing GFP+ tumor cells. Scale bar = 2mm. **E.** Longitudinal µCT imaging of mice. Yellow dotted lines indicate the boundaries of the orthotopic inguinal #4 mammary fat pad identified from whole mouse imaging of sh*Scr* and **F.** sh*Sox5* residual lesions. White arrow-heads indicate calcified areas within the orthotopic inguinal #4 mammary glands. **G.** µCT and PET in sh*Scr* and **H.** sh*Sox5* D48 residual lesions 75 minutes after injection of [$^{18}$F]NaF. Arrowheads indicate calcified areas in excised mammary glands. Orange dotted lines indicate lymph nodes.
